# Structural basis of drug recognition by the multidrug transporter ABCG2

**DOI:** 10.1101/2021.02.18.431786

**Authors:** Julia Kowal, Dongchun Ni, Scott M Jackson, Ioannis Manolaridis, Henning Stahlberg, Kaspar P Locher

## Abstract

ABCG2 is an ATP-binding cassette (ABC) transporter whose function affects the pharmacokinetics of drugs and contributes to multidrug resistance of cancer cells. While its interaction with the endogenous substrate estrone-3-sulfate (E_1_S) has been elucidated at a structural level, the recognition and recruitment of exogenous compounds is not understood at sufficiently high resolution. Here we present three cryo-EM structures of nanodisc-reconstituted, human ABCG2 bound to anticancer drugs tariquidar, topotecan and mitoxantrone. To enable structural insight at high resolution, we used Fab fragments of the ABCG2-specific monoclonal antibody 5D3, which binds to the external side of the transporter but does not interfere with drug-induced stimulation of ATPase activity. We observed that the binding pocket of ABCG2 can accommodate a single tariquidar molecule in a C-shaped conformation, similar to one of the two tariquidar molecules bound to ABCB1, where tariquidar acts as an inhibitor. We also found single copies of topotecan and mitoxantrone bound between key phenylalanine residues. Mutagenesis experiments confirmed the functional importance of two residues in the binding pocket, F439 and N436. Using 3D variability analyses, we found a correlation between substrate binding and reduced dynamics of the nucleotide binding domains (NBDs), suggesting a structural explanation for drug-induced ATPase stimulation. Our findings provide insight into how ABCG2 differentiates between inhibitors and substrates and may guide a rational design of new modulators and substrates.

## INTRODUCTION

ABCG2, also known as breast cancer resistance protein (BCRP), is an ATP-binding cassette (ABC) transporter expressed in many tissues and tissue barriers, including the blood–brain, blood–testis and maternal–fetal barriers[1–5]. It transports endogenous substrates such as uric acid and estrone-3-sulfate (E_1_S) and also removes exogenous, cytotoxic compounds from cells[6]. Like other multidrug ABC transporters, such as ABCB1 (P-glycoprotein) and ABCC1 (MRP1), ABCG2 is overexpressed in certain cancer cells, which contributes to drug resistance and affects chemotherapeutic intervention[7]. ABCG2 also affects the oral uptake rate and pharmacokinetics of many drugs[8]. Both ABCG2 and ABCB1 are expressed at the blood-brain barrier[9], where they prevent the entry of a broad variety of xenobiotics, limiting drug delivery to the brain[10–12].

ABCG2 recognizes and actively transports chemically distinct but mostly hydrophobic compounds, many of which are are polycyclic and have a relatively flat shape [13–17]. Transport substrates include topoisomerase inhibitors, tyrosine kinase inhibitors, and antimetabolites[16]. There is a substantial overlap between compounds recognized by ABCG2 and other multidrug ABC transporters. Intriguingly, there are also compounds that are substrates of one, but inhibitors of another multidrug ABC transporter[18]. A key example is tariquidar, a strong inhibitor of ABCB1 that was reported to be transported by ABCG2, albeit at a low rate [19–25].

Over the past few years, high-resolution structures of human ABCG2 have provided insight into its architecture, its substrate-binding pockets, and the conformational changes associated with ATP binding [26–28]. The structure of ABCG2 with its physiological substrate E_1_S bound in the central, cytoplasm-facing binding pocket helped identify key residues contributing to substrate recognition [28]. A comparison with structures of ABCG2 bound to inhibitory compounds [27] suggested that the binding sites of substrates and inhibitors partially overlap.

A recent study reported structures of ABCG2 in complex with the anti-cancer drugs imatinib, mitoxantrone, and SN38 at ∼4 Å resolution [29]. However, it remained unclear which ABCG2 residues are involved in specific drug-ABCG2 interactions and how larger drugs can be accommodated and transported by ABCG2. To address these questions, we determined three single-particle cryo-EM structures of nanodisc-reconstituted, substrate-bound ABCG2 in a pre-translocation state. We selected tariquidar, topotecan, and mitoxantrone for our studies. Tariquidar (647 Da) is an anthranilamide derivative that was originally developed as an ABCB1 inhibitor[19–24]. Topotecan (421.5 Da) is a topoisomerase I inhibitor that is in use to treat ovarian, cervical and small-cell lung cancer[30]. Mitoxantrone (444.5 Da) is a topoisomerase II inhibitor that is used to treat leukemia and multiple sclerosis [31]. To achieve sufficiently high resolution in the transmembrane domains of ABCG2 and visualize how substrates are bound, we investigated complexes of ABCG2 with Fab fragments of the specific antibody 5D3 that binds to the external surface of the transporter. This resulted in an increase of the particle mass and hence of the resolution of the obtained cryo-EM maps. Despite inhibiting substrate transport, the presence of 5D3-Fab does not interfere with substrate binding, as demonstrated by increased ATPase rates in the presence of the three drug compounds.

Our high-resolution structures combined with functional studies reveal how drugs are accommodated in the binding pocket of ABCG2 and which residues contribute to drug recognition. We demonstrate that substrates are not immobilized as firmly as inhibitors, but instead appear to be shifting in the binding cavity. By analyzing the structural variability between the single particles contributing to the 3D reconstructions, we found that the structural flexibility of the nucleotide binding domains (NBDs) is reduced by substrate binding although the substrate retains some mobility inside the drug-binding pocket. Our results provide a basis for understanding drug-ABCG2 interactions and for the future development of modulators and inhibitors.

## RESULTS

### *In vitro* characterization of ABCG2 function

We first reconstituted ABCG2 in liposomes and performed ATPase assays in the presence of the selected substrate compounds. We observed a concentration-dependent stimulation of the ATPase activity by topotecan, mitoxantrone and tariquidar (**Fig. 1**), which is in agreement with previous reports[13, 15–17, 25, 32]. The half-maximal effective concentration values (EC_50_) of substrate-induced ATPase stimulation were used to guide the substrate concentrations used for cryo-EM sample preparations. The EC_50_ values of mitoxantrone- and topotecan-induced ATP stimulation of ABCG2 were 18.3 µM and 11.7 µM, respectively, which is three orders of magnitude greater than the EC_50_ observed for tariquidar (0.08 µM) (**Table S1**), but in the same range as the EC_50_ of E_1_S-induced ATPase stimulation of ABCG2 alone or in the presence of 5D3-Fab [28].

**Fig. 1.**
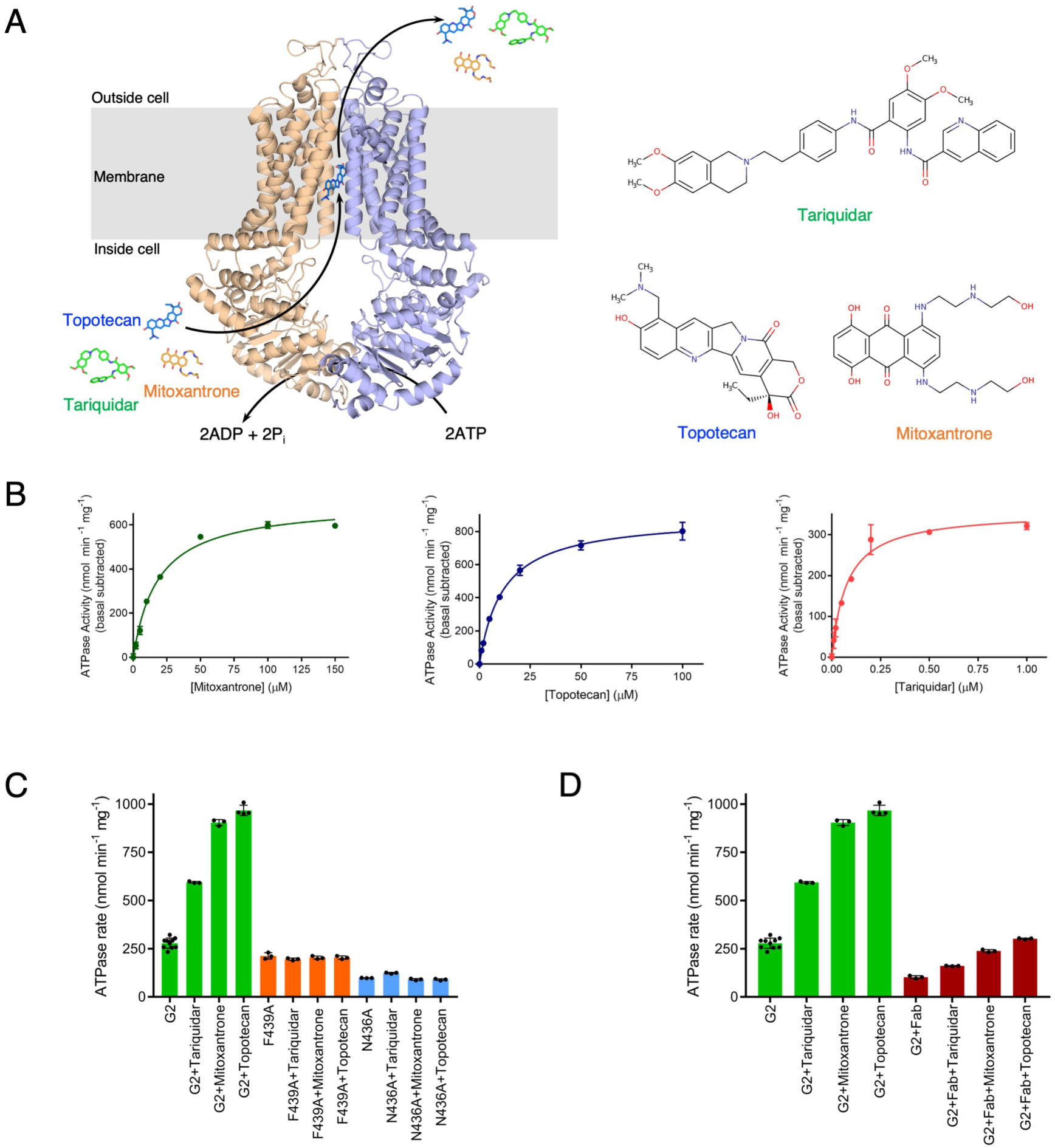
Substrate transport of the ABCG2. (A) Cartoon representation of ABCG2 with bound topotecan. ABCG2 monomers are colored salmon and blue. The substrates topotecan, mitoxantrone and tariquidar are shown as blue, orange and green sticks, respectively. The substrate transport cycle is indicated by arrows. Bound 5D3-Fab is omitted for clarity. Chemical structures of the substrates are shown on the right. (B) Determination of the EC_50_ of ATPase stimulation of ABCG2 in proteoliposomes by mitoxantrone, topotecan or tariquidar. The basal ATPase activity was normalized to zero. Each point represents the mean rate derived from technical replicates, for mitoxantrone and tariquidar n = 3, for topotecan n = 4. (C) ATPase activity of liposome-reconstituted wild-type and mutant ABCG2 (F439A, N436A) in the presence and absence of 0.5 μM tariquidar, 100 μM mitoxantrone, or 50 μM topotecan. The bars show means; error bars show standard deviations; and dots show rates derived from each technical replicate (same batch of liposomes). (D) Effect of 5D3-Fab on ATPase stimulation of liposome-stimulated ABCG2. ATPase rates were determined in the presence or absence of 5D3-Fab and either 0 or 0.5 μM tariquidar, 0 or 100 μM mitoxantrone, and 0 or 50 μM topotecan. Bars show means and dots show the rates derived from each technical replicate (same batch of liposomes). Error bars show the standard deviation.

We subsequently investigated ABCG2 variants carrying the mutations F439A and N436A, since these amino acids are involved in substrate binding[28]. The ATPase rates of these variant proteins were reduced, and they showed no stimulation by the drugs studied, indicating that interactions between the substrates and the side chain of F439 (π-stacking between phenyl ring and polycyclic molecules) and N436 (hydrogen bond formation) are essential for substrate binding (**Fig. 1C**). For our structural studies, we used 5D3-Fab to increase particle mass, stabilize the transporter and thus obtain higher resolution EM density maps. This antibody fragment binds to the extracellular side of ABCG2, inhibits its transport activity and slows down its ATPase rate[26]. To assess whether the antibody fragment interfered with drug binding to ABCG2, we measured the stimulation of ATPase activity of liposome-reconstituted ABCG2 by drugs in the presence and absence of 5D3-Fab. Our data suggests that 5D3-Fab does not interfere with the ATPase stimulation, and therefore drug binding to ABCG2 (**Fig. 1D** and **Table S2)**.

### Cryo-EM structures of drug-bound ABCG2

In order to obtain the single particle cryo-EM structures, we added 5D3-Fab to nanodisc-reconstituted ABCG2 and incubated the complex with either 100 μM topotecan, 150 μM mitoxantrone, or 1 μM tariquidar for 10 minutes at room temperature prior to plunge-freezing the grids. The cryo-EM data were processed with the Relion[33] and CryoSPARC[34] software suites. 3D classification revealed significant dynamics of the NBDs of ABCG2, with 35-70% of the particles containing separated and/or poorly resolved NBDs. After many rounds of 3D classifications, only the most stable conformations were selected and further refined to high-resolution (**Figs. S1-6, Table S3**).

The refined structures showed inward-facing conformations of ABCG2, with drugs bound in cavity 1 in a location similar to where the bound E_1_S was previously observed [26–28]. Although ABCG2 has two-fold symmetry only one copy of each substrate was observed (**Fig. 2A**). Furthermore, it would be impossible to accommodate two substrate molecules within the binding pocket, because their polycyclic structures would sterically clash. The transmembrane domains (TMDs), including the drug-binding pockets, were very well resolved (resolution higher than 3 Å), allowing fitting of the side chains of the key amino acids involved in substrate interactions. In all EM maps, we observed EM density features connecting substrates and ABCG2 side chains (**Fig. S7**). The strongest and most compact EM densities for substrates were observed for topotecan (3.39 Å) and mitoxantrone (3.51 Å). Intriguingly, despite tariquidar displaying the lowest EC_50_ of the three compounds, the EM density for the drug was less well-defined and more fragmented, suggesting that despite its higher affinity for ABCG2, tariquidar is either more flexible and/or bound in multiple orientations. Upon reprocessing of the data in CryoSPARC, we obtained a more compact and prominent density for tariquidar, albeit at the expense of a slightly lower overall resolution (3.5 Å) of the transporter (**Fig. 2, Fig. S7**). Both tariquidar-bound ABCG2 maps show that a submicromolar drug binds in a less defined way than other, weaker-binding drugs. Whereas the Relion map for ABCG2-tariquidar features the C-shaped tariquidar better, the CryoSPARC-generated map emphasizes the multiple binding modes.

**Fig. 2.**
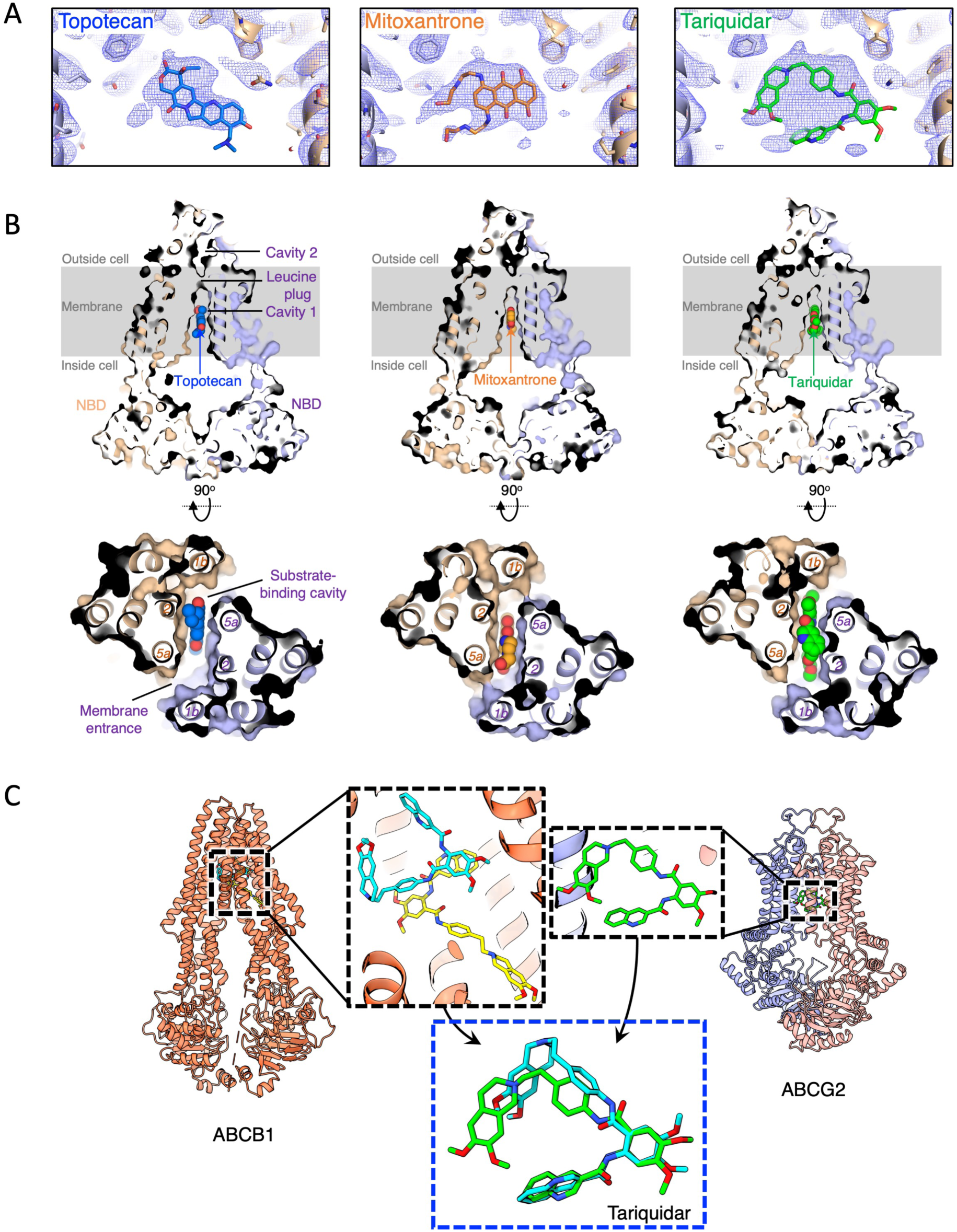
Drugs bound to ABCG2. (A) The binding pocket (cavity 1) of ABCG2 with drugs shown from the membrane plane. Non-symmetrized electron microscopy maps of ABCG2 with substrates shown as blue mesh. The map display levels were adjusted to show similar EM densities around the amino acids in the binding pocket in all maps. Bound topotecan, mitoxantrone and tariquidar molecules are shown as blue, orange and green sticks, respectively. (B) Fit of substrates in the slit-like binding cavity. Vertical slices through a surface representation of ABCG2 with bound substrates shown as blue (topotecan), orange (mitoxantrone) and green (tariquidar) spheres and labeled. Cavities 1 and 2, the leucine plug, as well as membrane and NBD regions are indicated. 5D3-Fab is omitted for clarity. Below, cavity 1 viewed from the cytoplasm, with NBDs of ABCG2 removed for clarity and with substrates. TM helices are labeled 1b, 2 and 5a. (C) Comparison of tariquidar molecule conformations from ABCB1- and ABCG2-tariquidar bound structures (**<u>PDB 7A6E</u>** from *Nosol et al*.[38] and **<u>PDB XXXX</u>** (this work), respectively). The two top zoomed-in panels show the close-up views of the binding pockets of tariquidar-bound ABCB1 (tariquidar molecules in U- and L-shape are colored cyan and yellow, respectively) and ABCG2 (tariquidar in C-shape, green sticks). Bottom panel (dashed, blue) shows the superposition of U- and C-shaped tariquidar molecules (green and cyan sticks) from both ABC-transporters.

### Substrate-ABCG2 interactions

All substrates bind in the same multidrug slit-like binding cavity (cavity 1) located at the level of the center of the membrane, below the leucine plug of ABCG2 and between TM helices 1b, 2 and 5a (**Fig. 2B**). The drug binding pocket lies on the two-fold symmetry axis of the ABCG2 dimer. In contrast, the Ko143-derived inhibitor MZ29 was found to bind in two copies shifted away from the two-fold symmetry axis, still almost entirely filling cavity 1, and another inhibitor, the tariquidar-derived MB136, was in one copy in a stretched conformation, occupying a larger space in the cavity compared to the substrates[27]. The shape of ABCG2’s binding pocket causes flat compounds to be preferred, in contrast to the globular molecules favored by ABCB1[35, 36] or ABCC1 transporters[37]. In this work we observed that the tariquidar molecule has to fold into a C-shape to fit into the cavity of ABCG2. Comparison of the conformations of the tariquidar molecule bound to the ABCG2 (this work, **Fig. 2A, Fig. S7**) and ABCB1 (from [38]) led to the conclusion that one of the tariquidar molecules that is lining the binding pocket of the ABCB1 is in a similar conformation as the ABCG2-bound tariquidar molecule (**Fig. 2C**).

The well-resolved and detailed maps of the three ABCG2-substrate structures allowed us to understand the specific requirements of substrate binding in cavity 1. Each substrate was polycyclic and bound between a pair of opposing phenyl rings of F439 by π-stacking interactions (**Fig. 3**). In addition, tariquidar and mitoxantrone formed hydrogen bonds with the side chain of N436, while for topotecan this interaction was absent because of the smaller size of the drug. For the hydrogen bonds the distance between the carboxamide group of N436 and the oxygen atom of the methoxy group of tariquidar is 2.9 Å and that between N436 and the hydroxyl-ethylamine tail of mitoxantrone is 3.2 Å. At the bottom of cavity 1, the presence of EM density linking the hydrophobic side chain of M549 and the substrates, suggests strong Van der Waals interactions and possibly an interaction between the sulfur of the methionine side chain and the edges of the aromatic ring systems of the drugs. Van der Waals interactions involve T542 (all substrates), V546 (mitoxantrone and tariquidar), and S440 and L405 (tariquidar) (for a full list see **Table 4**).

**Fig. 3.**
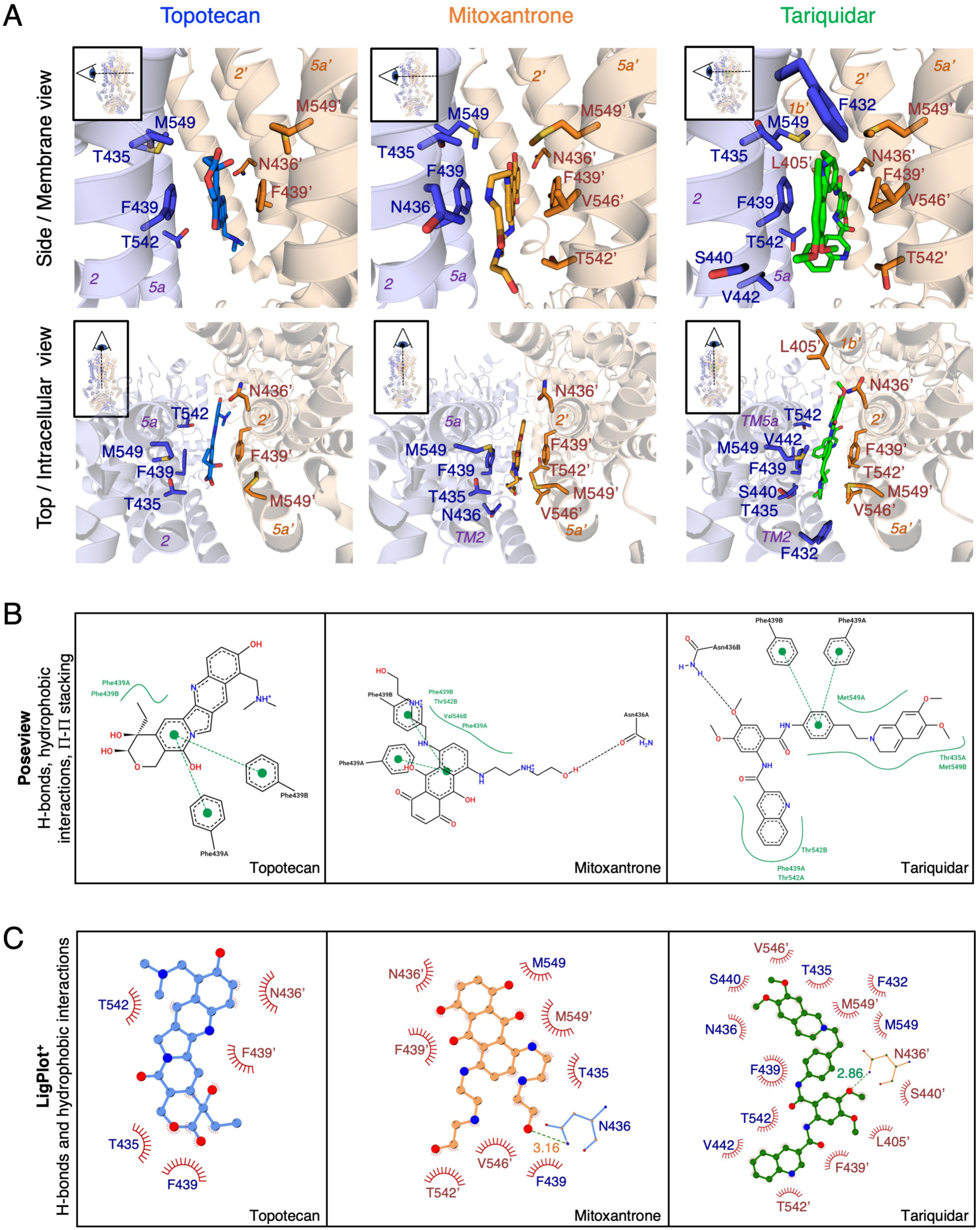
ABCG2-drug interactions. (A) Specific interactions between substrates topotecan (blue), mitoxantrone (orange) or tariquidar (green) and the ABCG2 residues in cavity 1, viewed from the membrane (top row) and from the extracellular site towards the cytoplasm (bottom row). Ribbon diagram of the ABCG2 homodimer, with individual ABCG2 monomers colored in blue and salmon. Interacting residues are shown as sticks and labeled. Substrates are shown as sticks. TM helices are labeled 1b’, 2, 2’, 5a and 5a’. (B) Two-dimensional visualizations of protein-substrate interactions generated from three-dimensional inputs. Hydrogen bonds are shown as black dashed lines, hydrophobic interactions as smooth contour lines between the respective amino acids and the substrate molecule, and π− π stacking as green dots connected by dashed line. Generated with Poseview(Stierand & Rarey, 2010). (C) Interactions between three substrates and ABCG2 side chains. Nonbonded interactions are represented by spoked red arcs and hydrogen bonds are indicated by dashed green lines. The amino acid residues are shown as single-letter abbreviations.

Our structures complement the data shown by Orlando and Liao[29]. In their work on ABCG2 in complex with chemotherapeutic drugs Orlando and Liao also observed density for bound drugs in cavity 1, although at lower resolution. While the general location of the drugs between the phenyl rings of F439 is similar, we find distinct orientations of bound mitoxantone and topotecan (depicted in **Fig. S8**). The mitoxantrone molecule in our structure is rotated almost 90° compared to the earlier study, and the topotecan molecule is flipped 180° to the structurally similar SN38 molecule in Orlando and Liao. Given our functional data, we can rule out a 5D3-Fab-induced artifact in our structures.

### Movement of drug substrates in the binding pocket

We investigated the conformational flexibility of drug-bound ABCG2 by performing 3D variability analyses[34]. This allowed the visualization of movements of the substrates inside the drug-binding cavity. The key observation was that the substrates kept sliding inside the slit-like cavity (**Movies S1-3**). We also noted structural differences akin to vibrations in the NBDs. Intriguingly, we found a correlation between the density of the NBDs and the observed densities of the substrates in between the TMDs. A larger separation of the NBDs resulted in less-resolved EM density of the substrate in cavity 1. Conversely, when the NBDs are closer together, the substrate density for is better defined. The mobility of the NBDs was particularly visible during 3D classifications, but a gradual selection of the most stable 3D class allowed refinement of the most stable conformation to high-resolution (**Figs. S2, S4, S6**). We conclude that when substrate molecules access the binding pocket the NBDs become ready to hydrolyze ATP and facilitate transport of the molecule (**Fig. 5**). Upon comparing our EM maps, the most “mobile” NBDs and the least-well defined drug density were observed for the ABCG2-tariquidar structure. Tariquidar was reported to adopt at least three different conformations[39]. Both our Relion and CryoSPARC EM density maps (**Fig. S7**) suggest that tariquidar can adopt a couple of distinct conformations similar to C-/U-conformation (**Fig. 2C, Fig. S9**).

## DISCUSSION

Among the endogenous substrates of ABCG2 are uric acid (168 Da) and estrone-3-sulfate (E_1_S, 350 Da). Both are small, heterocyclic compounds that contain two or four aromatic rings, respectively. The slit-like substrate binding cavity in ABCG2 is well-suited to recruit these two compounds and sandwich them between the phenyl groups of the F439 side chains. We found that despite the larger sizes and more complex structures, tariquidar, topotecan and mitoxantrone, interact with a similar set of amino acids in ABCG2 as the endogenous substrates[28]. However, the larger the exogenous compound, the greater are its interactions with distant amino acids, which are not observed for endogenous substrates. For example, residue T542 does not interact with E_1_S but does with all tested anti-cancer drugs (**Fig. 3**). The second largest substrate, mitoxantrone, has an additional hydrophobic interaction with V546 that is absent for topotecan. Finally, the largest drug, tariquidar (647 Da), has to alter its conformation to fit in the ABCG2 binding pocket and apart from the above-mentioned interactions observed for two previous substrates, it interacts with hydrophobic (V442, F432, L405) and polar (S440) amino acids located below the central part of the binding pocket.

The recently published structure of tariquidar-bound ABCB1 showed two molecules of tariquidar bound to the transporter [38]. The first adopted a globular, U-shaped (also referred to as C-shaped) conformation and is completely inside the binding pocket of ABCB1. The second tariquidar molecule has an L-shaped conformation and is located between the binding pocket and the access tunnel, where it interferes with conformational changes and thus inhibits ABCB1. The tariquidar molecule in ABCG2 does not adopt the elongated L-shaped conformation. It adopts a more compact C-shaped conformation similar to the tariquidar bound in the central binding pocket of ABCB1 (**Fig. 2C**). It had been proposed that the U-shaped tariquidar bound to ABCB1 could be transported if no second tariquidar molecule was present. In analogy, tariquidar bound to ABCG2 may represent a conformation of a transport substrate rather than an inhibitor. Consistent with this idea, it was previously shown that derivatives of tariquidar such as MB136[27], UR-ME22-1[40] or HM30181 analogues[21], developed as inhibitors of ABCG2, bind in an extended conformation rather than in a compact, C-shaped conformation. MB136 thereby interacts with the residues V401 and L405’ from the opposing ABCG2 monomers[27] (**Fig. 5C**). This suggests that the inhibitory characteristics of MB136 are associated with its extended conformation and interactions with the sides of cavity 1, since the substrate-like characteristics of tariquidar are associated with the fact that it could assume a coiled conformation, when lodged in cavity 1. Unlike ABCG2 inhibitors, which act as wedges to prevent ATP-induced NBD dimerization, ABCG2 substrates are compatible with the proposed peristaltic extrusion mechanism through the center of the transporter[28].

ABCG2 transports tariquidar, albeit at a very low rate[25]. We noticed that despite displaying the highest affinity (EC_50_ for ATPase stimulation in the sub-micromolar range) of the three drugs, tariquidar had the least well-defined EM density (**Figs. 2 and S7**), suggesting the presence of multiple conformations of tariquidar in the binding pocket of ABCG2. Importantly, while the size of tariquidar is larger than the typical ABCG2 substrate, its ability to adopt a C-shaped conformation makes it more compact and therefore a (slowly transported) substrate. If the classification of compounds into inhibitors or substrates is a continuum, tariquidar is an example of a compound around the middle of this continuum. This molecular interpretation could be useful in guiding the design of larger inhibitors that do not adopt a C-shape like tariquidar but maintain a set of interactions with residues on the TM1b helices as observed in tariquidar derivatives[41].

By performing 3D variability analyses, we obtained insight into the dynamics of ABCG2 during substrate binding. The density features representing bound substrates among a continuous family of generated 3D structures for all substrates (**Movies S1-3**) allowed us to conclude that the substrate (i) alters the tilt of TM1 narrowing the entrance to the cavity, and (ii) correlates with an ordering of the NBDs, structurally explaining the observed ATPase stimulation in the presence of drugs (**Fig. 4**). We hypothesize that similar effects may take place in other ABC-transporters. Both topotecan and mitoxantrone intercalate between two DNA bases at the active sites of topoisomerases I and II, such that the enzymes are trapped in the cleavage complex bound to the DNA. When comparing the interactions of these anticancer drugs either with DNA or F439 residues in the ABCG2 binding pocket (**Fig. S10**), we observed that the distance between base pairs and the drugs is ∼0.5 Å smaller than between the phenylalanine residues and the drugs. This can rationalize the much weaker interactions between drug substrates and ABCG2, which allows the molecules to move within the binding pocket and, upon ATP binding to ABCG2, get transported.

**Fig. 4.**
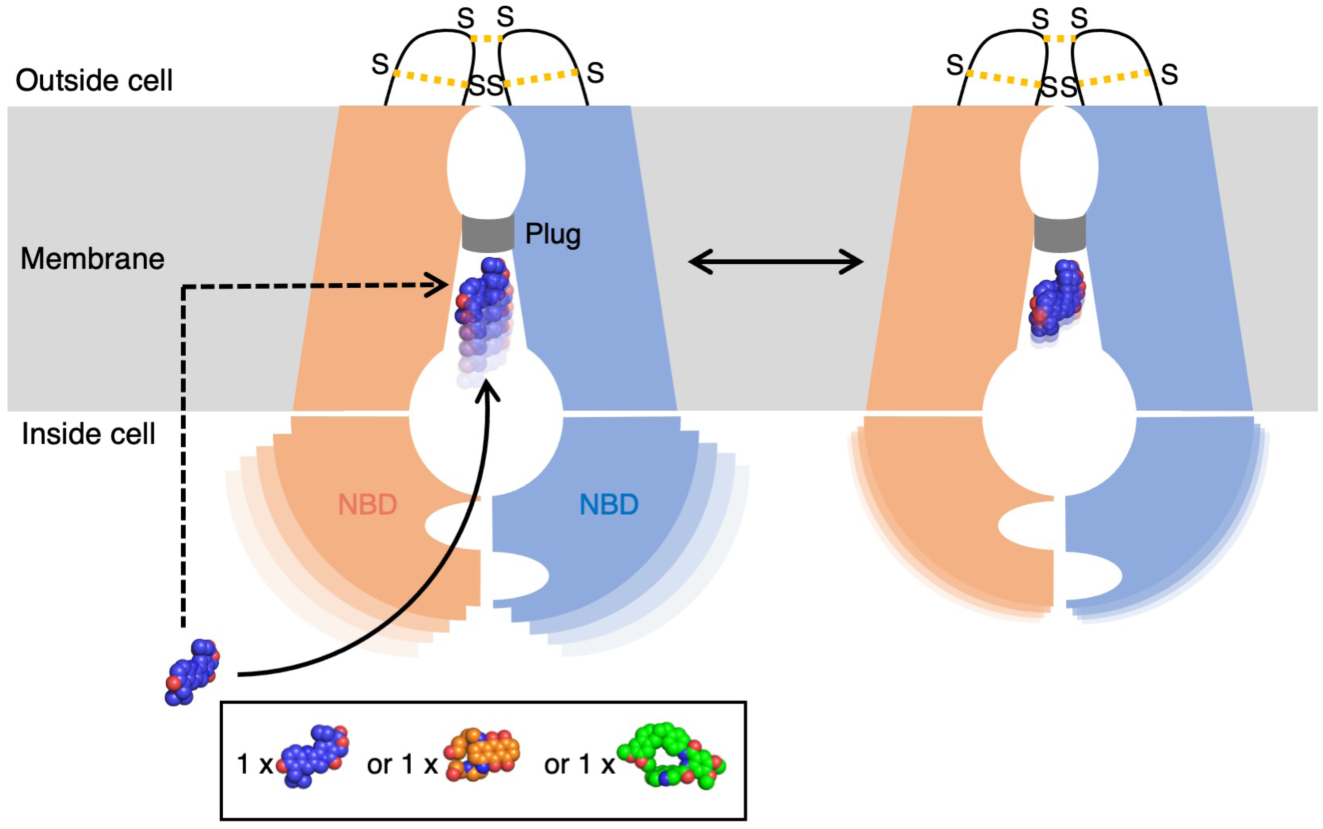
Effect of substrate binding on the NBD structure. Schematic of ABCG2 binding the substrate molecule. The binding of single substrate molecule (blue spheres) in the center of cavity 1 is a continuous process which requires a lot of movement. Substrate molecule is neither static nor locked in cavity 1. ABCG2 monomers are colored salmon and blue, disulfide bridges located at EL3 are indicated by yellow dashed lines connecting sulfur (S) atoms, and the leucine plug is shown as a gray bar between the cavities.

**Fig. 5.**
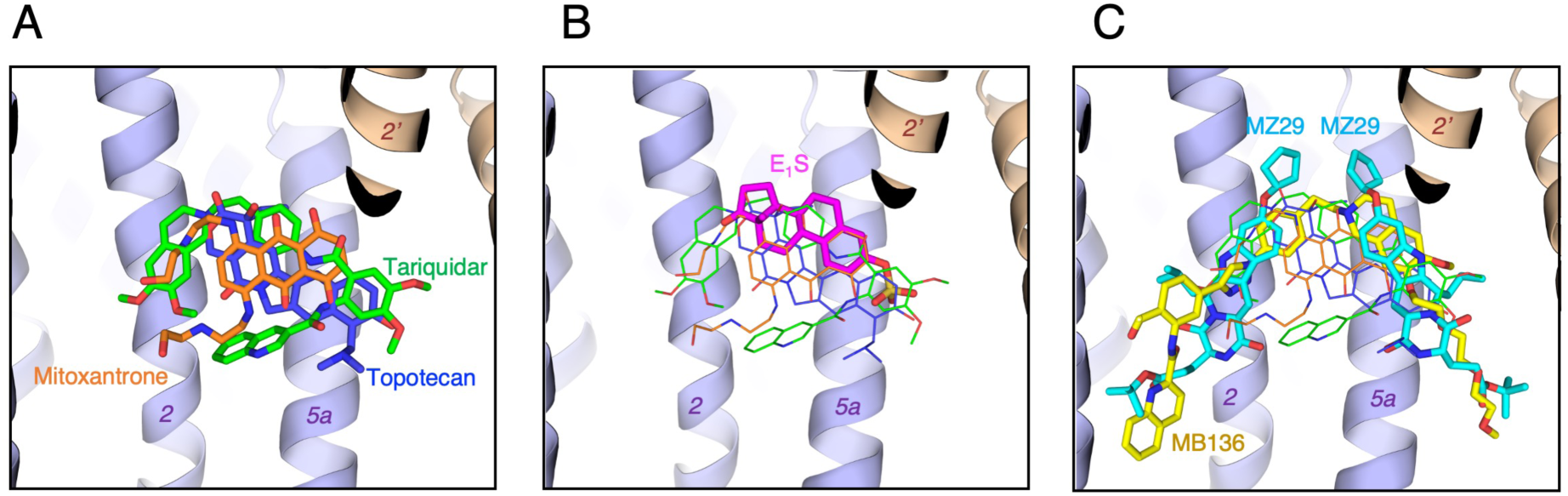
Substrates versus inhibitors in cavity 1 of ABCG2. Overlay of bound ligands from different ABCG2 structures after superposition of the TMDs. TM helices are labeled 2, 2’ and 5a. (A) Overlay of the three substrates topotecan, mitoxantrone and tariquidar (this work) shown as blue, orange and green sticks, respectively. (B) Substrates from (A) shown as thin sticks and estrone 3-sulfate (E_1_S) (thick, magenta sticks, from **<u>PDB 6HCO</u>**). (C) Overlay of substrates from (A) shown as thin sticks and ABCG2 inhibitors MZ29 (cyan sticks, from **<u>PDB 6ETI</u>**) and MB136 (yellow sticks, from **<u>PDB 6FEQ</u>**).

The presented ABCG2-drug structures explain the structural basis of substrate binding in ABCG2. However, despite the recent progress, further improvements in obtaining high resolution and well-defined ligand densities as well as increased structure determination throughput would be required to make thorough structure-based drug design campaigns viable. Even at this advanced stage of structural studies, we conclude that total stabilization of a substrate molecule in the ABCG2 binding pocket is questionable. The understanding of the substrate’s requirements and how they differ from those of inhibitory compounds might make the approaches of designing better ABCG2 inhibitors more successful.

## MATERIALS AND METHODS

### Expression and purification of wild-type human ABCG2 and 5D3-Fab

Human ABCG2, containing an amino (N)-terminal Flag tag, was expressed in HEK293-EBNA (Thermo Fisher Scientific) cells and purified as described previously[26–28].

5D3 hybridoma cells, producing the 5D3 monoclonal antibody, were obtained from B. Sorrentino. The cells were cultured in WHEATON CELLine Bioreactors, according to the manufacturer’s protocol, and 5D3-Fab was then purified from the supernatant, as described in the Fab Preparation Kit protocol (Thermo Fisher Scientific).

### Nanodisc preparation of ABCG2

The membrane scaffold protein (MSP) 1D1 was expressed and purified and ABCG2 was reconstituted into brain polar lipid (BPL)/cholesterol hemisuccinate (CHS) nanodiscs as described previously[26–28]. To generate the ABCG2-Substrate samples for cryo-EM studies, ABCG2 was first mixed with a threefold molar excess of 5D3-Fab before reconstitution. After size-exclusion chromatography (SEC) using a Superdex 200 10/300 column (GE Healthcare), the ABCG2-Fab complex was incubated with 100 μM topotecan, 150 μM mitoxantrone or 1 μM tariquidar for 10 min at room temperature before cryo-EM grids preparation.

### ABCG2-liposome preparation

A BPL/cholesterol (BPL/chol) (Avanti Polar Lipids) mixture was prepared at a 4:1 (w/w) ratio as described previously[42]. Briefly, the BPL/chol mixture was extruded through a 400-nm polycarbonate filter and destabilized with 0.17% (v/v) Triton X100. Detergent-purified ABCG2 was then mixed with BPL/chol at a 100:1 (w/w) lipid/protein ratio. Detergent was removed with BioBeads, and proteoliposomes were spun at 100,000g, resuspended in buffer 25 mM HEPES, pH 7.5, 150 mM NaCl at a final lipid concentration of 10 mg/ml, and the reconstitution efficiency was determined[43].

### ATPase assays and determination of the EC_50_ of mitoxantrone, topotecan or tariquidar stimulation

ATP-hydrolysis activity was measured using a previously described technique[44]. Proteoliposome samples were pre-incubated with the specified concentrations of mitoxantrone, topotecan or tariquidar for 10 min at 37 °C prior to the addition of 2 mM ATP and 10 mM MgCl_2_ to start the hydrolysis reaction. To assess the effect of 5D3-Fab, ABCG2 proteoliposomes were freeze-thawed five times in the presence of a three-fold molar excess of 5D3-Fab to ensure that 5D3-Fab was inside the proteoliposomes before extrusion[26, 27]. Data were recorded at four-time intervals (0, 5, 15 and 30 min) and subsequent ATPase rates were determined using linear regression in GraphPad Prism 7.00. Rates were corrected for the orientation of ABCG2 in proteoliposomes[26]. To determine the EC_50_ of mitoxantrone, topotecan and tariquidar stimulation, we plotted the ATPase rates against the substrate concentration, and generated curves using the nonlinear regression Michaelis–Menten analysis tool in GraphPad Prism 7.00.

### Sample preparation and cryo-EM data acquisition

All cryo-EM grids of wild type ABCG2 with topotecan, mitoxantrone or tariquidar were prepared with Vitrobot mark IV (FEI) with the environmental chamber set to 100% humidity and 4°C. Aliquots of 4 μL of purified ABCG2-Fab complexes with substrates at a protein concentration of approximately 0.5 mg/mL were placed onto Quantifoil carbon grids (R1.2/1.3, 300 mesh, copper) that had been glow-discharged for 45 seconds at 25 mA using Pelco easiGlow 91000 Glow Discharge Cleaning System. Grids were blotted for 1.5-2.0 s and flash-frozen in a mixture of liquid ethane and propane cooled by liquid nitrogen.

The final data set of ABCG2-tariquidar-Fab was composed of 5894 super-resolution movies. The ABCG2-tariquidar-Fab grids were imaged with a Titan Krios (FEI) electron microscope operated at 300 kV, equipped with a Gatan K3 direct electron detector and Gatan Quantum-LS energy filter (GIF), with a slit width of 20 eV to remove inelastically scattered electrons. Movies were recorded semi-automatically with EPU software (Thermo Fisher Co.), in super-resolution counting mode with a defocus range of –0.4 to –2.5 μm and a super-resolution pixel size of 0.33 Å/pixel. Movies were 1.0 s long, dose-fractionated into 40 frames, with a frame exposure of 2.0 e^−^/Å^2^. All stacks were gain-normalized, motion-corrected, dose-weighted and then binned 2-fold with MotionCor2[45]. The defocus values were estimated on the non-dose-weighted micrographs with Gctf[46].

The ABCG2-topotecan-Fab dataset was composed of 3403 movies. The movies were collected on Titan Krios microscope (300kV), equipped with a Gatan Quantum-LS energy filter (20 eV zero loss filtering) on a K2 Summit detector with the automation of SerialEM[47]. Movies were recorded in counting mode with a physical pixel size of 0.82 Å/pixel, in a defocus range from −0.8 to −2.8 μm, with a frame exposure of 1.55 e^−^/Å^2^ and 40 frames for each movie (8s exposure). All the movie stacks were gain-normalized, motion-corrected and dose-weighted with the program MotionCor2[45].

The ABCG2-mitoxantrone-Fab data was composed of 6192 movies in total. The movie data was collected on a Titan Krios microscope (300kV) with an automation software SerialEM[47], equipped with a Gatan Quantum-LS energy filter (20 eV zero loss filtering). The movies were recorded with a K2 Summit direct electron detector (counting mode) with a pixel size of 0.64 Å/pixel. Movies were recorded in a defocus range from −0.6 to −2.5 μm, with a frame exposure rate of 1.50 e^−^/Å^2^ (40 frames for each movie, 8s in total). All the movie stacks were gain-normalized, motion-corrected and dose-weighted with the program MotionCor2[45]. The micrographs were sorted according to different parameters such as: Iciness, Sample Drift, Defocus, Astigmatism and Resolution of CTF Fit (in Focus[48]), in order to remove the bad images.

### Image processing

Image processing of the ABCG2-tariquidar-Fab complex was performed both in Relion 3.1[33] and CryoSPARC2[34]. From the 5,894 selected dose-weighted micrographs, a total of 1,326,686 particles were picked within Relion. After five rounds of 2D classifications and selection, we obtained a final set of 737,525 particles which were 3D classified within 3 classes using as reference a low-passed initial model obtained from the small dataset processed in CryoSPARC2 (*ab-initio* reconstruction). The particles from the best 3D class, containing 644,542 (87.4%) of particles, were re-extracted (384 x 384 box) without binning, and 3D refined with soft mask covering the full density of the complex. The resolution of this map was 3.62 Å. Subsequently, the series of 3D refinements including CTF refinement with anisotropic magnification correction and beam-tilt correction, per-particle CTF refinement, Bayesian polishing and refinement with a soft mask without detergent belt and constant domain of Fab were performed and yielded the 3.27 Å resolution map. To further improve the map resolution, the particles were applied for another 3D classification without alignments, without a mask. At this point we noticed that only 27% (166,727) of the particles contributed to the 3D class with stable NBDs. This class was further 3D refined in the Relion with an angular sampling rate of 0.5 degree and local searches set to 1.8 degrees, and with C1 or C2 symmetry applied (maps at 3.12 Å and 2.98 Å resolution, respectively, with automatically determined B-factors of −74 Å^2^ and −77.5 Å^2^). In order to get clearer density for the tariquidar molecule the focused refinement with mask covering transmembrane (TM) region only was performed. The resulting masked map at 3.58 Å had better defined density of the drug, which was in a C-shape conformation.

In parallel, the same set of particles was processed in CryoSPARC v2.14 where 2D classification, *ab-initio* 3D reconstruction and homogeneous 3D refinement with C1 symmetry were performed. The resulting 3D map at 3.5 Å resolution (with B-factor adjusted to −100 Å^2^) showed more compact density for the drug molecule in comparison to Relion’s maps. Further, using the set of particles contributing to the final 3D volume, the 3D variability analyses were performed in CryoSPARC v2.14.

All of the aligned movies of the ABCG2-topotecan-Fab and ABCG2-mitoxantrone-Fab were prepared in Focus[48] and then imported into CryoSPARC2[34]. The CTF and defocus values were estimated on the dose-weighted micrographs. To generate a template for particles picking for the ABCG2-topotecan-Fab sample, 3000 particles were picked manually and 2D classified. The second round of particle picking was performed using the templates and resulted in 775,993 particles. After two rounds of 2D classifications 417,949 particles were selected. A 3D template was created using the *ab-initio* 3D reconstruction (with C1 symmetry, similarity 0.1). The initial 3D classification was followed by 3D refinement of the best 3D class containing 203,789 particles. The resulted map with C1 symmetry had an overall resolution of 3.50 Å. Because of the possible heterogeneity within the ABCG2-topotecan-Fab sample another round of *ab-initio* 3D reconstruction (similarity 0.65) plus 3D hetero-refinement with three classes were performed and the final 3D class contained 133,832 particles. The final subset was subjected to series of 3D refinement jobs (Homogeneous Refinement > Non-uniform Refinement > Local Refinement) applying C1 and C2 symmetries (maps at 3.39 Å and 3.14 Å resolution, respectively).

For ABCG2-mitoxantrone sample, a similar processing was performed as described above. However, the main problem which we were facing during the processing, was the misalignment of the particles resulting in an averaged fake density. Therefore, we performed two independent *ab-initio* 3D reconstruction jobs, with the similarity parameter of 0.1 for C2 map, in order to achieve better resolution, and of 0.65 for C1 map to determine the substrate density located at the two-fold symmetry axis of the transporter, followed by the heterogenous 3D refinements. Final refinements resulted in EM density maps at 3.51 Å and 3.35 Å resolution, for C1- and C2-symmetrized maps, respectively.

The particle stacks contributing into best 3D maps of ABCG2 with drugs topotecan, mitoxantrone and tariquidar were used for further 3D Variability Analysis, which is a tool in CryoSPARC v2 for exploring both discrete and continuous heterogeneity in single particle cryo-EM data sets. In order to remove the signal coming from the micelle the masks without detergent belt made in Chimera were used. The simple linear “movies” of volumes were generated for 3 eigenvectors. Outputs were low-pass filtered during optimization to 4 Å and visualized with 3D Variability Display tool in CryoSPARC v2.

### Model building and refinement

For the generation of an initial model of human ABCG2-tariquidar-Fab, we used a post-processed non-symmetrized map from Relion at an overall resolution of 3.12 Å. Guided by the structure of ABCG2-MB136-Fab (**<u>PDB: 6FEQ</u>**)[27], we performed manual building and fitting using the software Coot[49]. The EM density was of excellent quality in the transmembrane region and variable domain of the antibody region and allowed the unambiguous building of ABCG2. In the NBD region, we carried out manual fitting and modifications where the resolution allowed. Subsequently, the manual fitting of the tariquidar molecule into the density was performed. For the final fit of the tariquidar molecule the Relion’s map from the TM region-focused refinement and the 3.5 Å map generated in CryoSPARC2 were used. Geometry restraints for tariquidar, cholesterol and phospholipid molecules were generated in eLBOW[50]. The complete model was refined against the working map in PHENIX[51] using real space refinement.

For the ABCG2-topotecan and -mitoxantrone structures model building, we used the previous ABCG2_EQ_-estrone-3-sulfate structure (**<u>PDB 6HCO</u>**) as an initial template. The PDB files were fitted into the resolved EM map low-passed to 6 Å in Chimera. Model building was performed manually in Coot. Geometry restraints for mitoxantrone and topotecan molecules were downloaded from PDB. After rebuilding, the atomic model was refined in PHENIX[51] using the option real space refinement with a standard parameters for geometry minimization including the global real-space refinement, non-crystallographic symmetry (NCS) and secondary structure, Ramachandran plot and rotamer restraints. The complete models were refined against the working maps in PHENIX[51] using real space refinement.

### Figure preparation

Figures were prepared using the programs PyMOL (PyMOL Molecular Graphics System, DeLano Scientific), ChimeraX and GraphPad Prism 7.00 (GraphPad Software).

## Abbreviations

ABC: ATP-binding cassette
E_1_S: estrone-3-sulfate
cryo-EM: cryo-electron microscopy
BCRP: breast cancer resistance protein
NBDs: nucleotide binding domains
TMDs: transmembrane domains

## Data availability

Atomic coordinates for ABCG2–topotecan-Fab, ABCG2–mitoxantrone-Fab and ABCG2–tariquidar-Fab (including only the variable domain of 5D3-Fab) were deposited in the Protein Data Bank under accession codes **<u>PDB XXXX, PDB YYYY</u>** and **<u>PDB ZZZZ</u>**, respectively. Electron microscopy data for the three structures were deposited in the Electron Microscopy Data Bank under accession codes **<u>EMD-XXXX</u>** (ABCG2–topotecan-Fab), **<u>EMD-YYYY</u>** (ABCG2–mitoxantrone-Fab) and **<u>EMD-ZZZZ</u>** (ABCG2–tariquidar-Fab).

## Acknowledgements

This work was in part supported by the Swiss National Science Foundation (NCCR TransCure). We thank the staff at the Scientific Center for Optical and Electron Microscopy (ScopeM, ETH Zurich, Switzerland) for support during cryo-EM data collection.

## Author contributions

I.M expressed and purified ABCG2 and 5D3-Fab. S.M.J performed ATPase activity assays and proteoliposomes experiments. S.M.J and I.M. reconstituted ABCG2 into liposomes and lipidic nanodiscs. J.K prepared all cryo-grids and with I.M. collected cryo-EM data for ABCG2-tariquidar-Fab. J.K determined the structure of ABCG2-tariquidar-Fab. D.N. and H.S collected cryo-EM data and determined the structures of ABCG2-topotecan-Fab and ABCG2-mitoxantrone-Fab. J.K. and K.P.L refined and validated the structure ABCG2-tariquidar-Fab. D.N. refined and validated the structures ABCG2-topotecan-Fab and ABCG2-mitoxantrone-Fab. K.P.L, J.K., S.M.J and I.M conceived the project. K.P.L, J.K., S.M.J and I.M planned the experiments. J.K. and K.P.L wrote the manuscript; all authors contributed to revisions.

## Competing financial interests

The authors declare no competing financial interests.

## SUPPLEMENTARY TABLES

**Table S1.** The EC_50_ of substrate ATPase stimulation determined using all curves from **Figure 1B** with the error of the fit (standard deviation) shown.

| Substrate | $EC_{50}$ ($\mu M$) (ATPase) |
| --- | --- |
| Mitoxantrone | $18.3 \pm 2.2$ |
| Topotecan | 11.7 ± 0.3 |
| Tariquidar | 0.08 ± 0.01 |

**Table S2.** The fold stimulation in the ATPase activity of ABCG2 caused by the addition of substrate in the presence and absence of Fab. Calculated from the data in **Figure 1D**.

| Substrate | G2 fold stimulation | G2+Fab fold stimulation |
| --- | --- | --- |
| + Tariquidar | 2.1 | 1.6 |
| + Mitoxantrone | 3.2 | 2.3 |
| + Topotecan | 3.5 | 2.9 |

**Table S3.** Cryo-EM data collection, refinement and validation statistics.

| Cryo-electron microscopy data collection and processing |  |  |  |
| --- | --- | --- | --- |
| Microscope | FEI Titan Krios | FEI Titan Krios | FEI Titan Krios |
| Voltage (keV) | 300 | 300 | 300 |
| Camera | Gatan K2-Summit | Gatan K2-Summit | Gatan K3 |
| Electron exposure (e<sup>-</sup>/Å<sup>2</sup>) | 1.55 | 1.5 | 2.0 |
| Energy filter slit width (eV) | 20 (Gatan Quantum-LS (GIF)) | 20 (Gatan Quantum-LS (GIF)) | 20 (Gatan Quantum-LS energy filter (GIF)) |
| Pixel size (Å) | 0.82 | 0.64 | 0.66 |
| Defocus range (μm) | (-0.8) - (-3.0) | (-0.6) - (-2.0) | (-0.4) - (-2.5) |
| Magnification (nominal) | 60'975x (165kx) | 78'125x (215kx) | 76'000 (130kx) |
| Number of frames per movie | 40 | 40 | 40 |
| Number of good micrographs | 3'403 | 6'192 | 5'894 |
| Initial particles | 775'993 | 614'891 | 1'326'686 |
| Final particles | 133'832 | 166'965 (91'543 for C1 map) | 166'727 |
| Symmetry imposed | C1,<br>/C2 (model building) | C1,<br>/C2 (model building) | C1 (model building),<br>/ C2 |
| Map resolution (Å) | 3.39 / 3.14 | 3.51 / 3.35 | 3.12 / 2.98 |
| FSC threshold | 0.143 | 0.143 | 0.143 |
| Map resolution range (Å) | 20 -2.70 | 20 -2.60 | 20-2.60 |
| EMDB ID | EMD-XXXX | EMD-XXXX | EMD-XXXX |
| **Refinement** |  |  |  |
| Non-hydrogen atoms | 12393 | 12394 | 12529 |
| Protein residues | 1582 | 1582 | 1582 |
| Ligands | TTC:1<br>NAG:2 | MIX:1<br>NAG:2 | TAR:1<br>NAG:2<br>CLR:3<br>PEE:1 |
| RMSD bonds (Å) | 0.006 | 0.005 | 0.009 |
| RMSD angles (°) | 0.723 | 0.757 | 0.752 |
| Ramachandran favored (%) | 90.63 | 90.56 | 92.11 |
| Ramachandran Allowed (%) | 9.37 | 9.31 | 7.89 |
| Ramachandran Outliers (%) | 0.00 | 0.13 | 0.00 |
| Rotamer outliers (%) | 0.30 | 0.00 | 5.56 |
| Cβ outliers (%) | 0.00 | 0.00 | 0.00 |
| All-atom clashscore | 12.62 | 20.23 | 15.39 |
| MolProbity Score | 2.14 | 2.33 | 2.75 |
| PDB ID | XXXX | XXXX | XXXX |

**Table S4.**
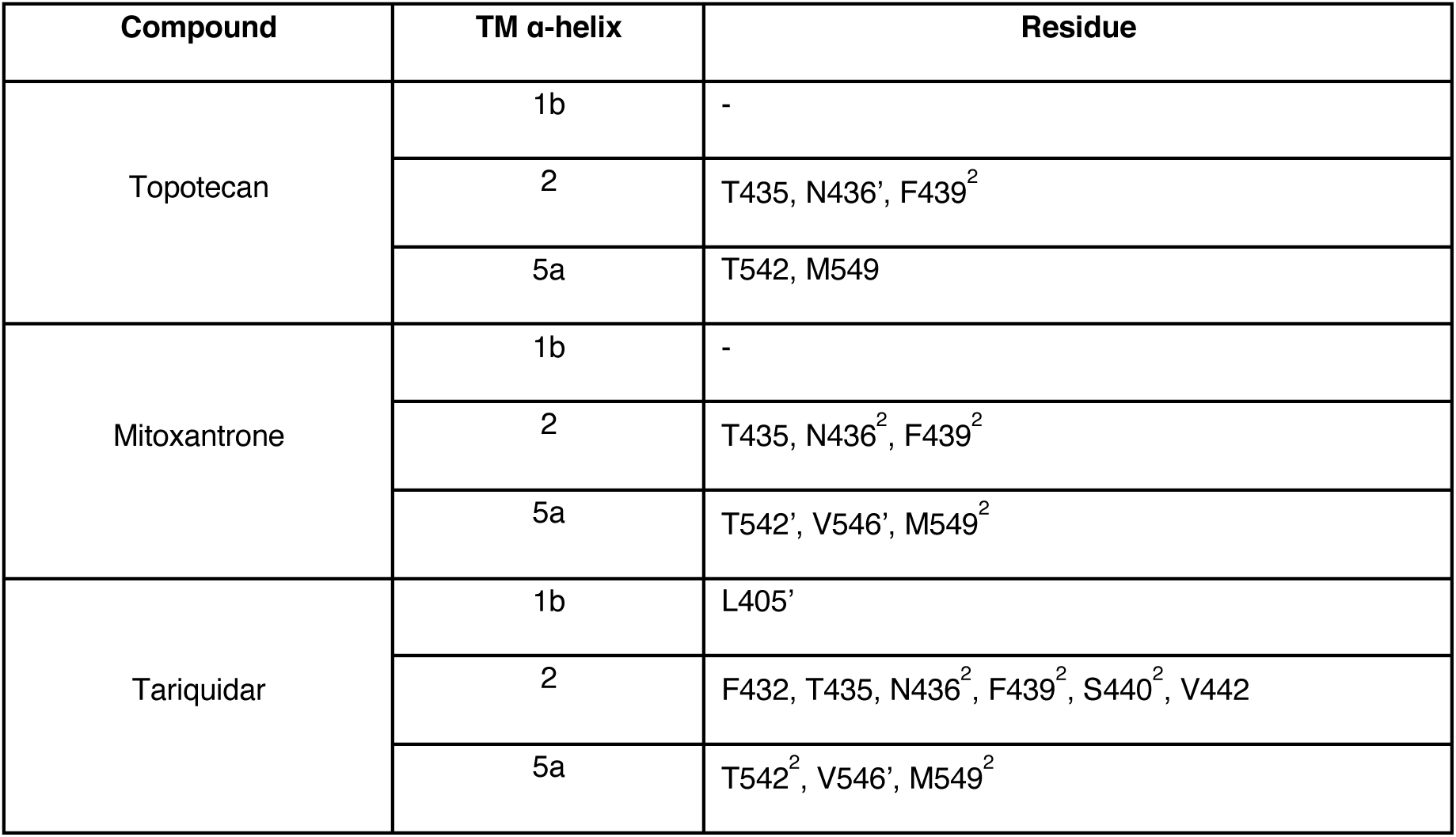
ABCG2 residues within 4 Å that interact with anticancer drugs in the ABCG2-substrate-Fab structures as shown on **Figure 3**. The prime (’) corresponds to the second half of the ABCG2, the (^2^) to both halves of transporter.

## SUPPLEMENTARY FIGURES

**Figure S1.**
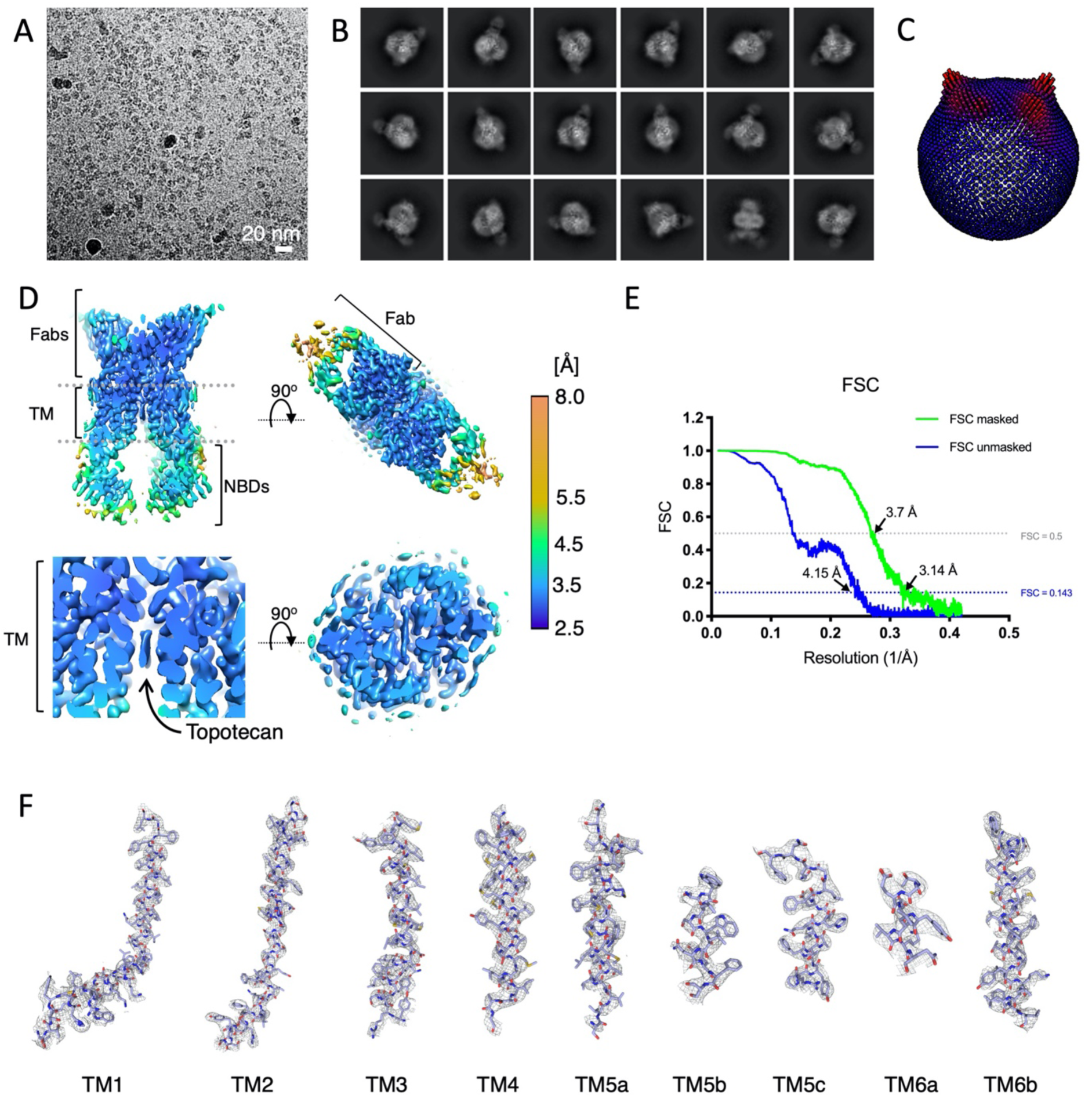
Cryo-EM map generation of ABCG2-topotecan-Fab and resolution estimation. **(A)** An example micrograph (drift-corrected, dose-weighted, and low-pass filtered) of the nanodisc-reconstituted ABCG2-topotecan-Fab. White scale bar, 20 nm. **(B)** Eighteen representative 2D class averages of the final round of 2D classification, sorted by decreasing order of the number of particles assigned to each class. **(C)** Angular distribution plot for the final reconstruction. **(D)** Full view of the final CryoSPARC B-factor-sharpened map of ABCG2-topotecan-Fab coloured by local resolution in Å, with the clipping plane in the middle of the molecule. On the top right corner is shown 5D3-Fabs region, in the bottom line the TM region with topotecan substrate density. **(E)** FSC from the CryoSPARC2 auto-refine procedure of the C2-symmetrized unmasked half-maps (blue) and the half-maps after masking (green). Horizontal dotted lines (blue and black) are drawn for the FSC = 0.143 and FSC = 0.5 criterion, respectively. For both the unmasked and the masked FSC curves, their intersection with the FSC = 0.143 and the FSC = 0.5 lines are marked by arrows, and the resolutions at these points are indicated. **(F)** Fitting of the all TM helices of ABCG2 in the EM density map. A region of up to 2 Å around the atoms is shown. TM helices are shown as sticks. Density is shown as a grey mesh.

**Figure S2.**
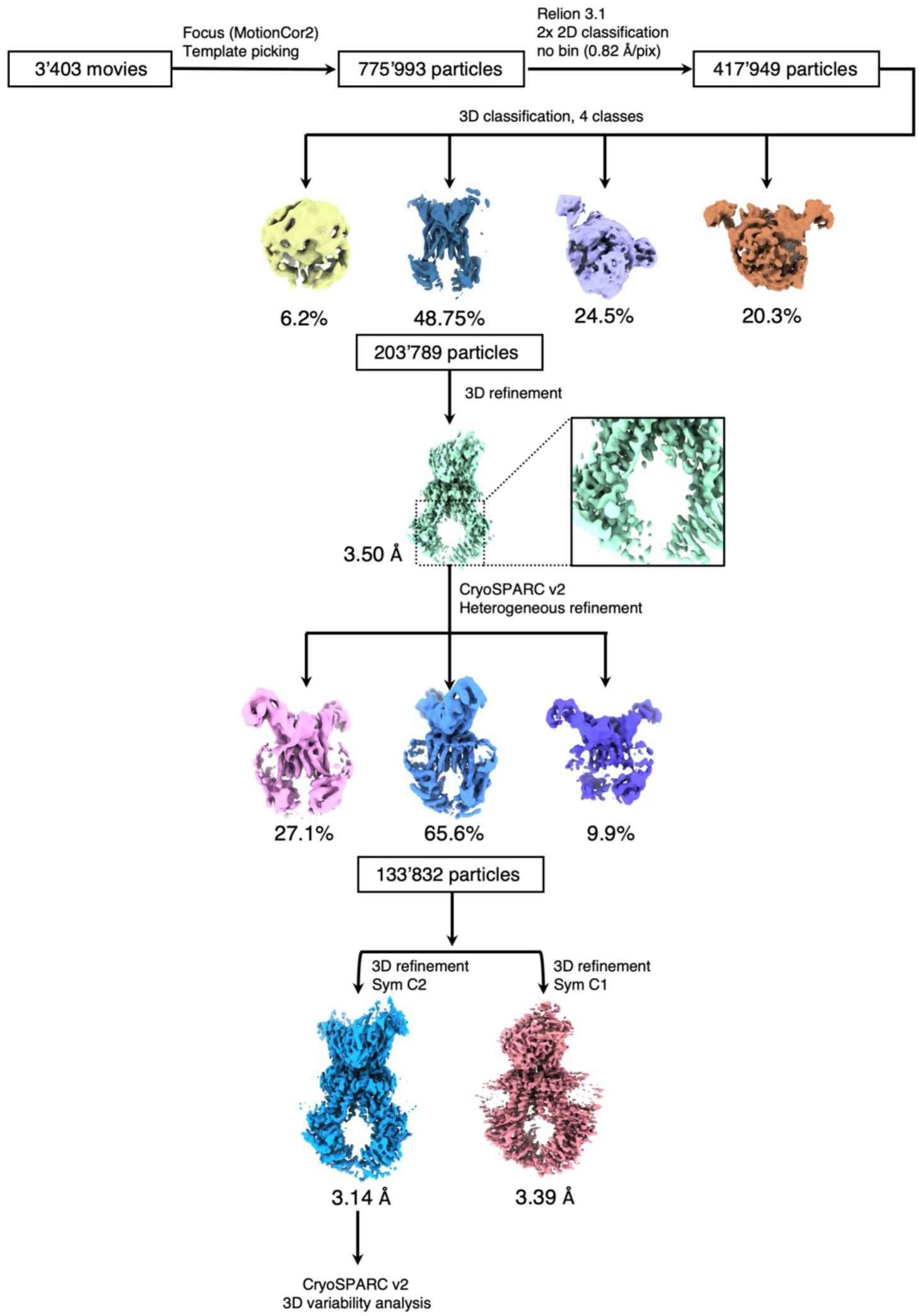
Flow chart for cryo-EM data processing and structure determination of the ABCG2-topotecan-Fab complex.

**Figure S3.**
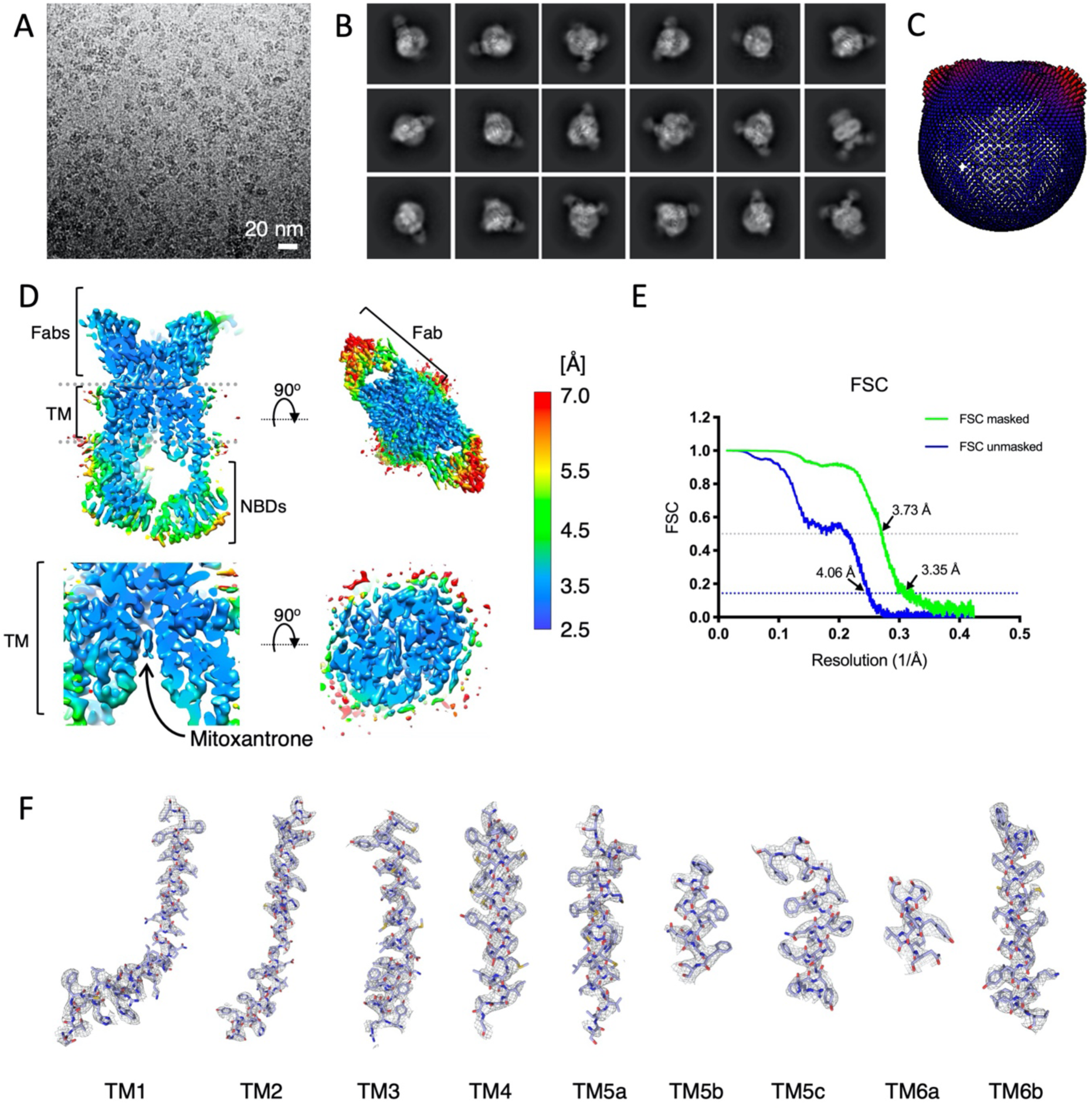
Cryo-EM map generation of ABCG2-mitoxantrone-Fab and resolution estimation. **(A)** An example micrograph (drift-corrected, dose-weighted, and low-pass filtered) of the nanodisc-reconstituted ABCG2-mitoxantrone-Fab. White scale bar, 20 nm. **(B)** Eighteen representative 2D class averages of the final round of 2D classification, sorted in decreasing order by the number of particles assigned to each class. **(C)** Angular distribution plot for the final reconstruction. **(D)** Full view of the final CryoSPARC B-factor-sharpened map of ABCG2-mitoxantrone-Fab coloured by local resolution in Å, with the clipping plane in the middle of the molecule. The 5D3-Fabs region is shown in the top right corner, bottom row - the TM region with the mitoxantrone substrate density. **(E)** FSC from the CryoSPARC2 auto-refine procedure of the C2-symmetrized unmasked half-maps (blue) and the half-maps after masking (green). Horizontal dotted lines (blue and black) are drawn for the FSC = 0.143 and FSC = 0.5 criterion, respectively. For both the unmasked and the masked FSC curves, their intersection with the FSC = 0.143 and the FSC = 0.5 lines are marked by arrows, and the resolutions at these points are indicated. **(F)** Fitting of all the TM helices of ABCG2 in the EM density map. A region of up to 2 Å around the atoms is shown. TM helices are shown as sticks. Density is shown as a grey mesh.

**Figure S4.**
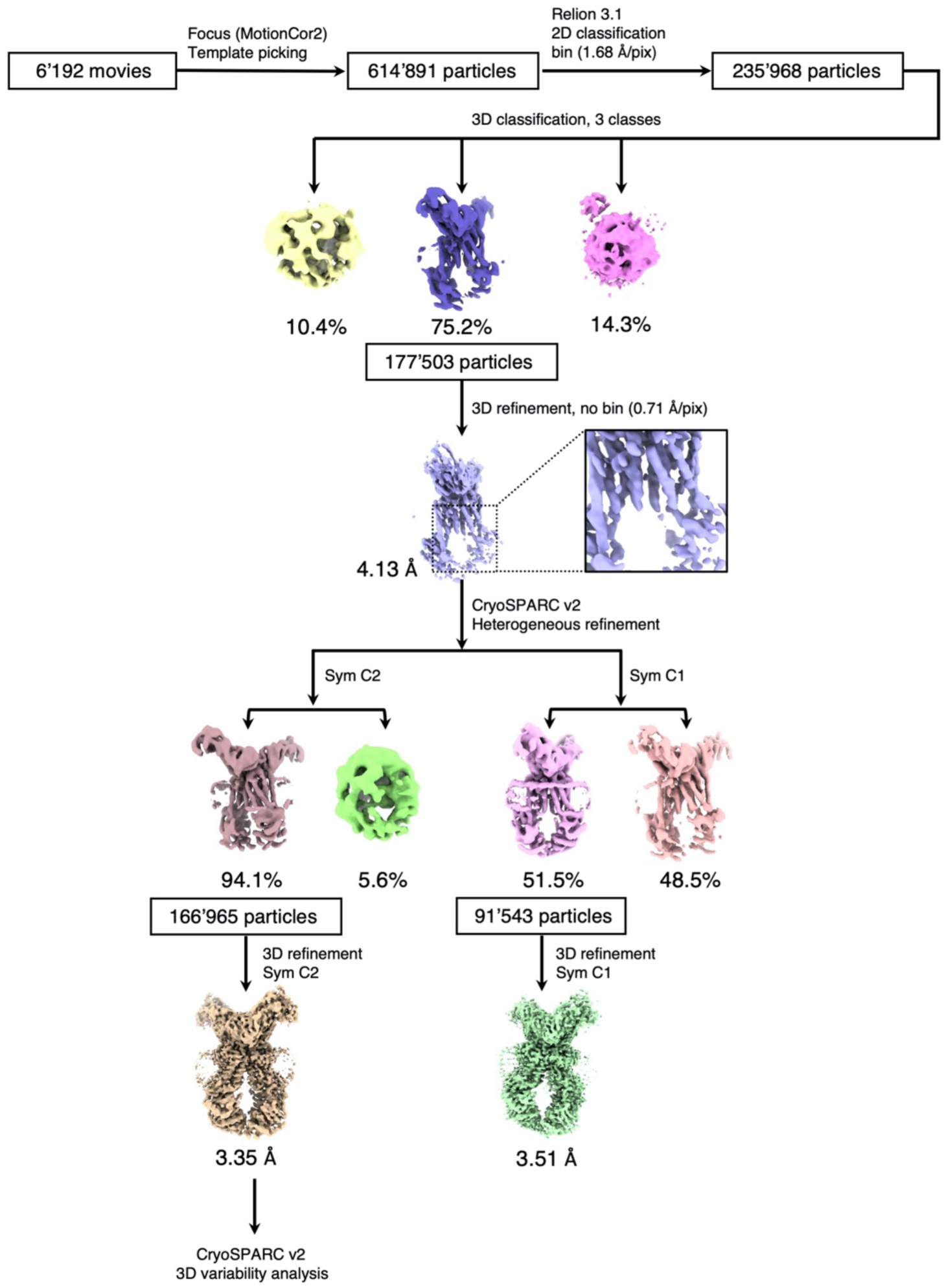
Flow chart for cryo-EM data processing and structure determination of the ABCG2-mitoxantrone-Fab complex.

**Figure S5.**
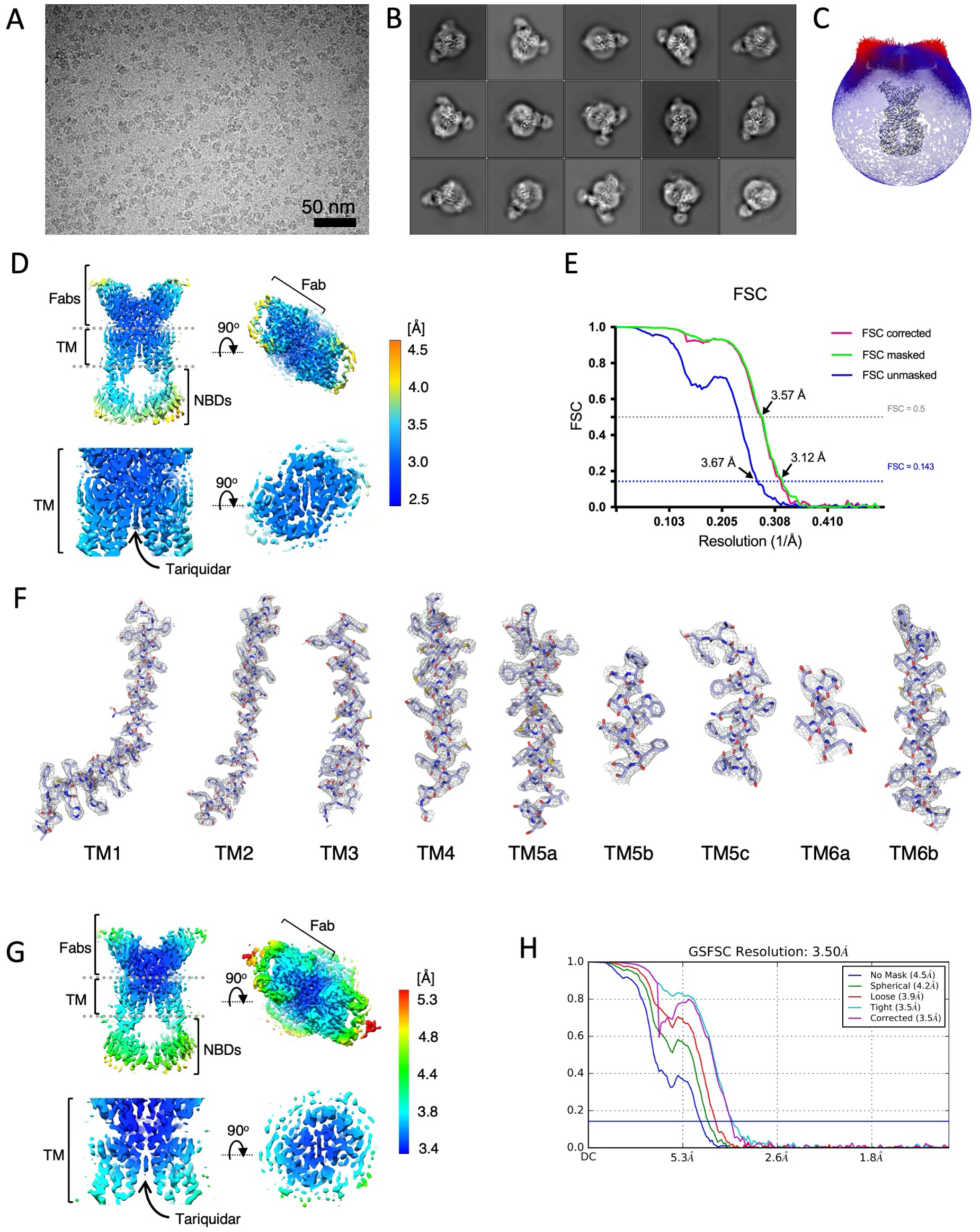
Cryo-EM map of ABCG2-tariquidar-Fab generation and resolution estimation. **(A)** An example micrograph (drift-corrected, dose-weighted, and low-pass filtered) of the nanodisc-reconstituted ABCG2-tariquidar-Fab. Black scale bar, 50 nm. **(B)** Fifteen representative 2D class averages of the final round of 2D classification, sorted in decreasing order by the number of particles assigned to each class. **(C)** Angular distribution plot for the final reconstruction. **(D)** Full view of the RELION local-resolution-filtered map of ABCG2-tariquidar-Fab, coloured by local resolution in Å as calculated in Relion 3.1, with the clipping plane in the middle of the molecule. In the top right corner is shown 5D3-Fabs region, bottom row - the TM region with tariquidar substrate density. **(E)** FSC from the RELION auto-refine procedure of the non-symmetrized unmasked half-maps (blue), the half-maps after masking (green), and the half-maps after masking and correction for the influence of the mask (pink). Horizontal dotted lines (blue and black) are drawn for the FSC = 0.143 and FSC = 0.5 criterion, respectively. For both the unmasked and the corrected FSC curves, their intersection with the FSC = 0.143 and the FSC = 0.5 lines are marked by arrows, and the resolutions at these points are indicated. **(F)** Fitting of the all TM helices of ABCG2 in the EM density map. A region of up to 2 Å around the atoms is shown. TM helices are shown as sticks. Density is shown as a grey mesh. **(G)** Full view of the local-resolution-filtered map of ABCG2-tariquidar-Fab from CryoSPARC, coloured as in (D). **(H)** FSC curves from the CryoSPARC 3D refinement. Horizontal blue line is drawn for the FSC = 0.143 criterion. The FSC curves calculated between two unmasked, masked or corrected half maps are shown.

**Figure S6.**
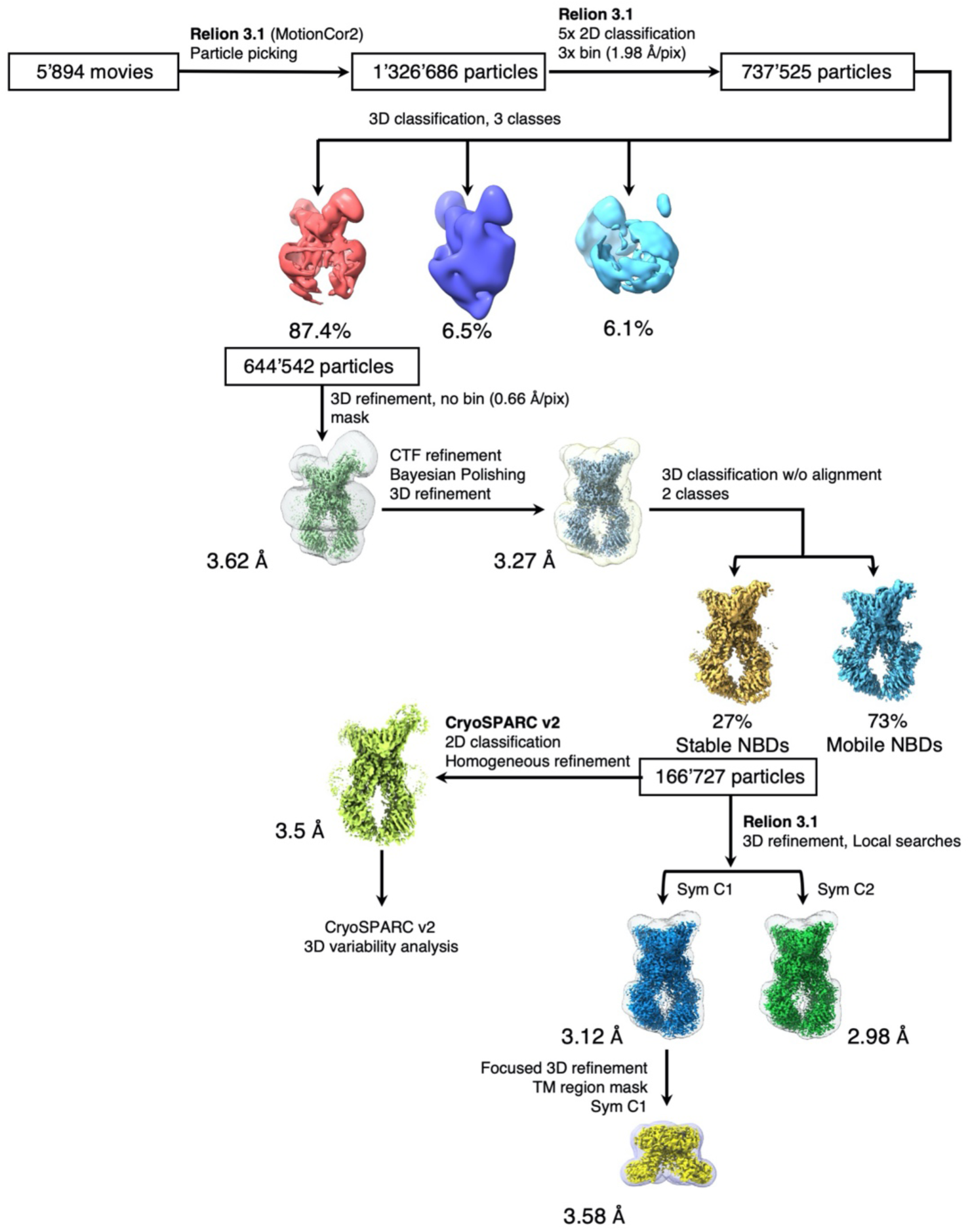
Flow chart for cryo-EM data processing and structure determination of the ABCG2-tariquidar-Fab complex.

**Figure S7.**
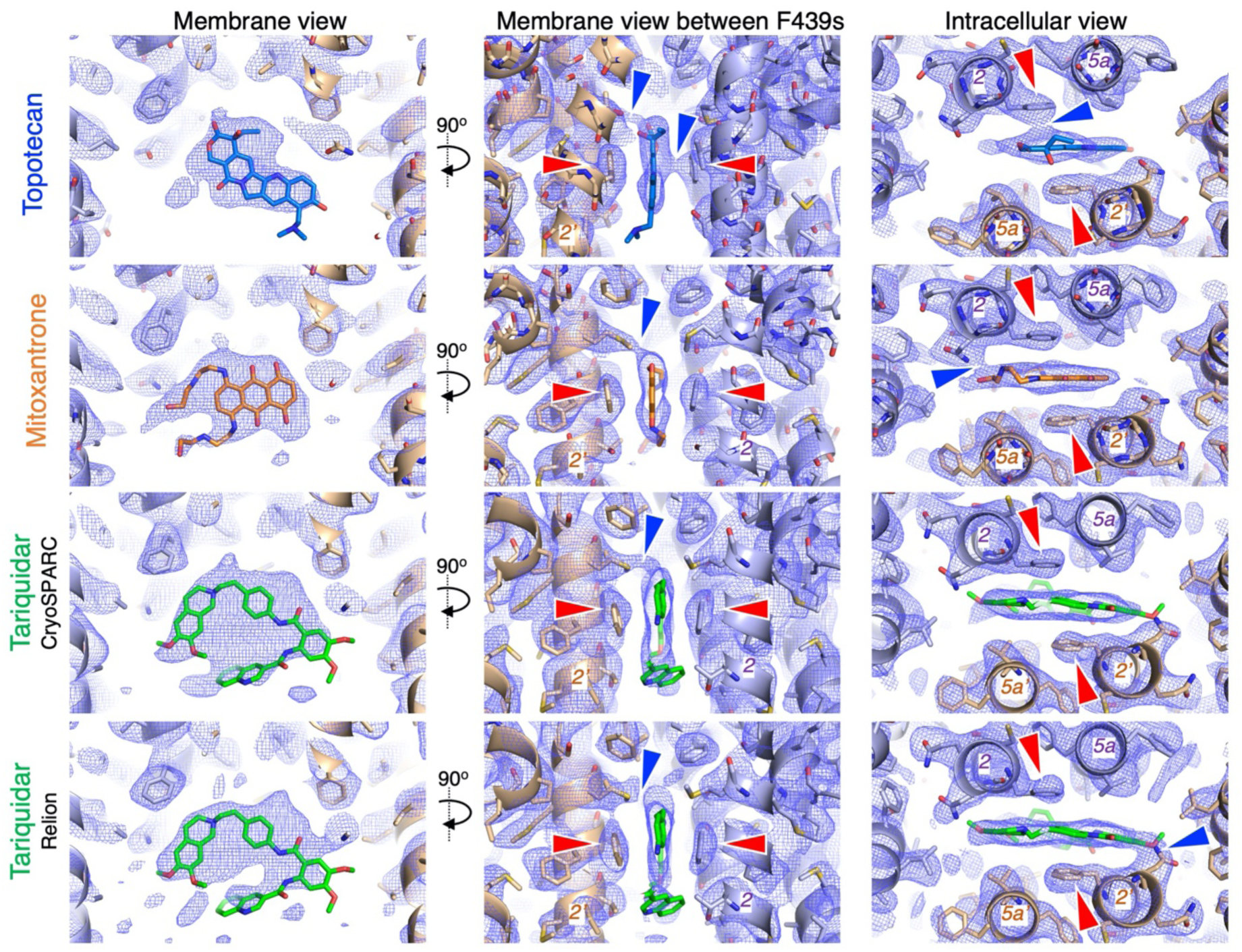
EM maps of ABCG2 with drug molecules. Non-symmetrized electron microscopy maps of ABCG2 with substrates are shown as blue mesh. The map display levels were adjusted to show similar EM densities around the amino acids in the binding pocket in all maps. Bound topotecan, mitoxantrone and tariquidar molecules are shown as blue, orange and green sticks, respectively. Side chains of F439 are marked with red arrows. Blue arrows mark sites where the EM density of the surrounding side chains and bound drugs are connected. TM helices are labeled 2, 2’, 5a and 5a’. Two maps, generated in CryoSPARC and Relion, of tariquidar-bound ABCG2 are shown.

**Figure S8.**
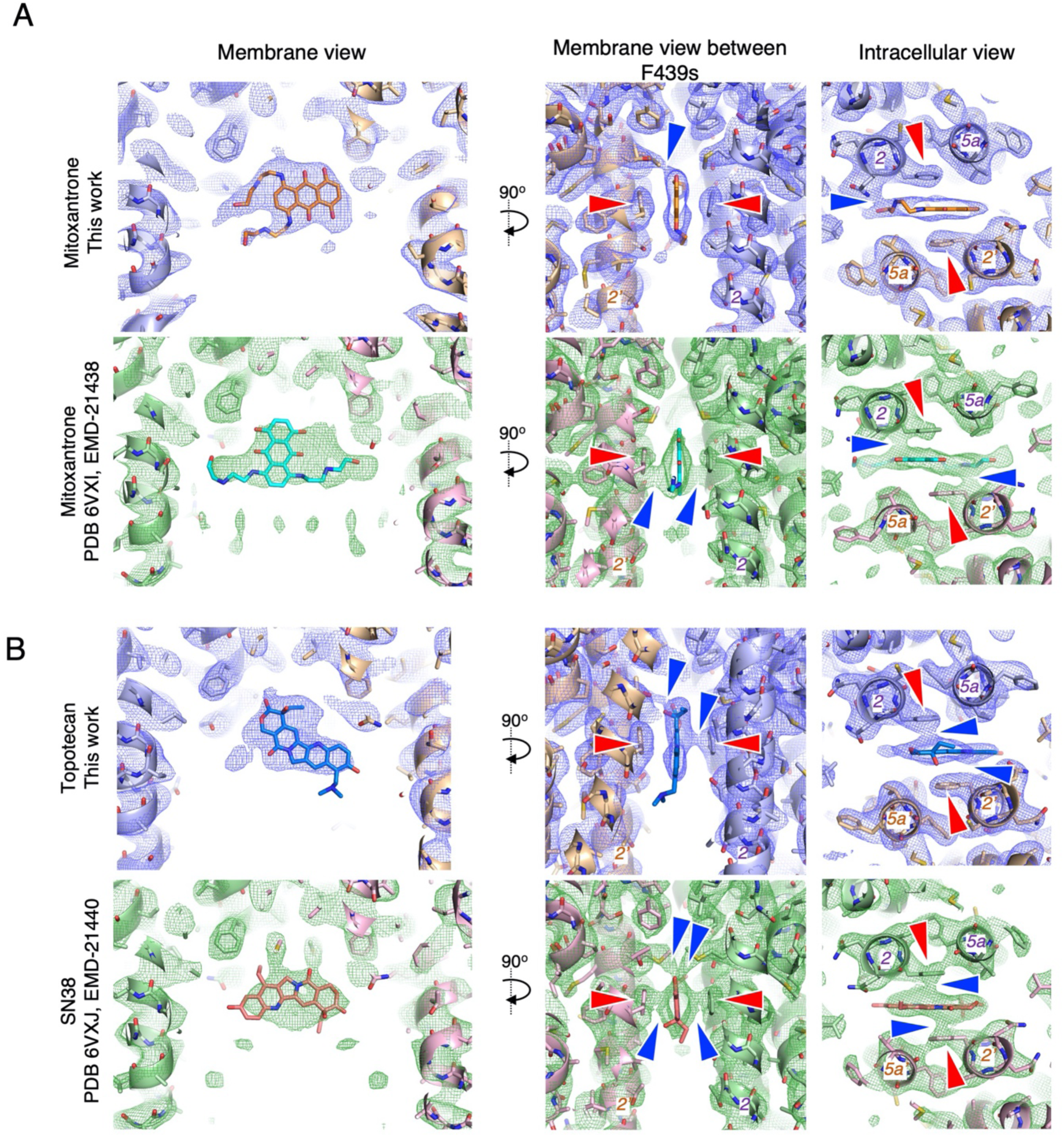
Comparison of the cryo-EM maps and models of ABCG2 with mitoxantrone, topotecan and SN38 anticancer drugs. Current ABCG2-mitoxantrone and ABCG2-topotecan models were fitted into the EM density maps (blue mesh) and compared with recently published maps/models of ABCG2-mitoxantrone and ABCG2-SN38 (green mesh). Two membrane views and an intracellular view are presented for each of the structure. **(A)** EM densities and models of ABCG2-mitoxantron. Top row, the model from our current work (mitoxantrone is orange). Bottom row, the ABCG2-mitoxantrone C2-symmetrized map from Orlando and Liao (**<u>PDB 6VXI, EMD-21438</u>**, mitoxantrone is cyan). the mitoxantrone presented in Orlando and Liao[29] is rotated almost 90° relative to our structure since its anthracene ring system is positioned vertically in cavity 1, while its long hydroxyl-ethylamine chains are horizontally stretched on both sides of cavity 1. In our ABCG2-mitoxantrone structure, the anthracene rings are oriented horizontally in the transporter and one of the hydroxyl-ethylamine tails forms a hydrogen bound with N436. The EM density is compact and does not suffer from C2-symmetrization effects. The map levels were readjusted to display similar EM densities around binding pocket of ABCG2. Levels 4σ and 6σ, as defined by Pymol, were used to display the EM maps (blue mesh – this work, green mesh – Orlando and Liao (2020) work), respectively. Labels as on the **Figure S7**. **(B)** EM densities and models of ABCG2-topotecan (this work, top row) and ABCG2-SN38 (**<u>PDB 6VXJ, EMD-21440</u>**, bottom row, molecule is pink). Labels/adjustments as in **(A)** and on **Figure S7**. SN38 molecule, whose chemical structure is similar to topotecan is located in the binding cavity upside down (180° rotation along the x axis) in comparison to our model.

**Figure S9.**
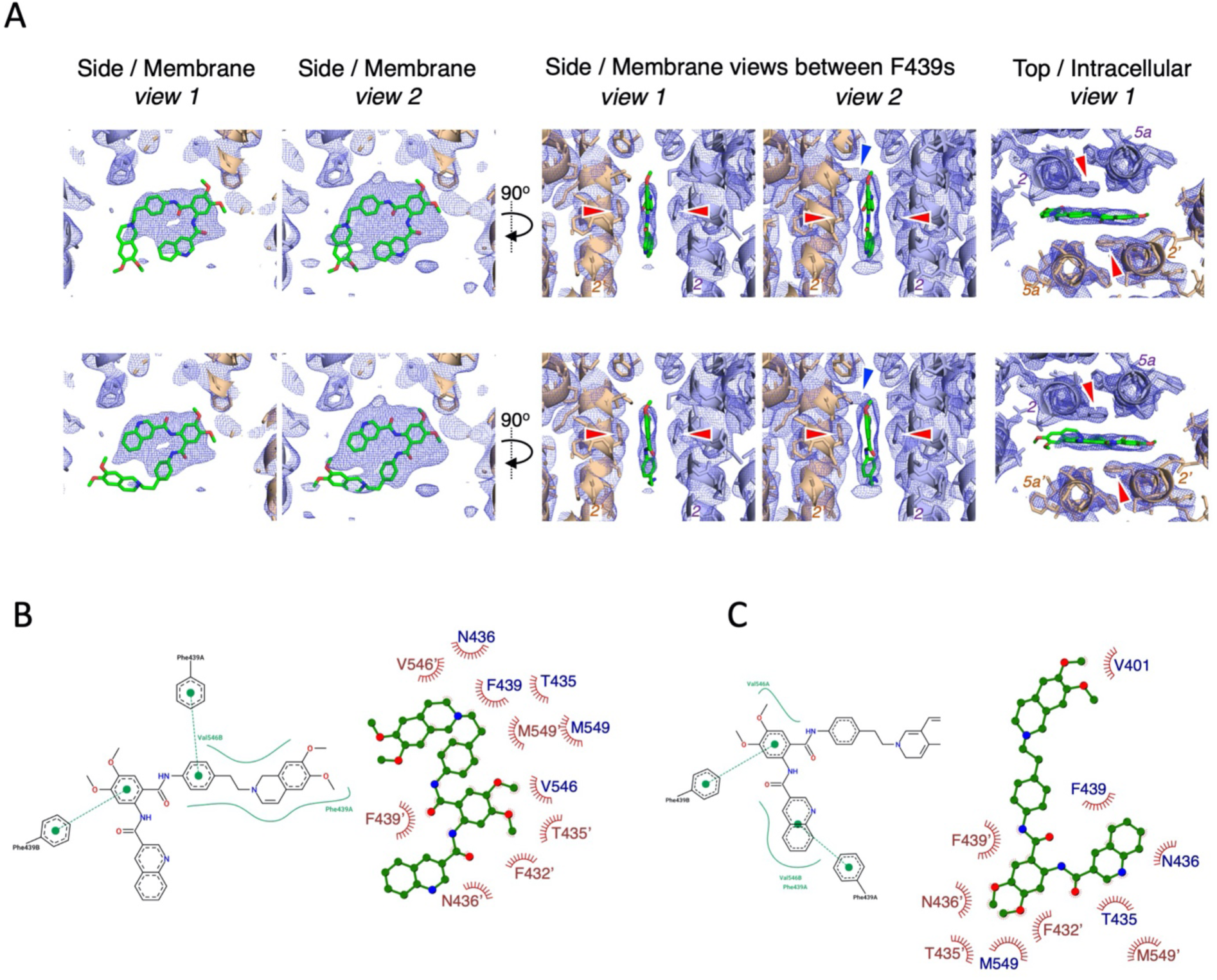
Two alternative conformations of tariquidar molecule fitted in ABCG2-tariquidar-Fab map obtained with CryoSPARC. **(A)** Two possible conformations of tariquidar fitted into the ABCG2-Tariquidar-Fab map. Tariquidar molecule shown as green sticks, map as blue mesh. Labels as in **Figure S7**. In *view 1* and *view 2* map levels of 9.0σ and 6.0σ, respectively, as defined by Pymol, were used to display the EM map. **(B)** Two-dimensional visualisations of protein-substrate interactions generated from three-dimensional input for both conformations from (A) with Poseview and LigPlot+.

**Figure S10.**
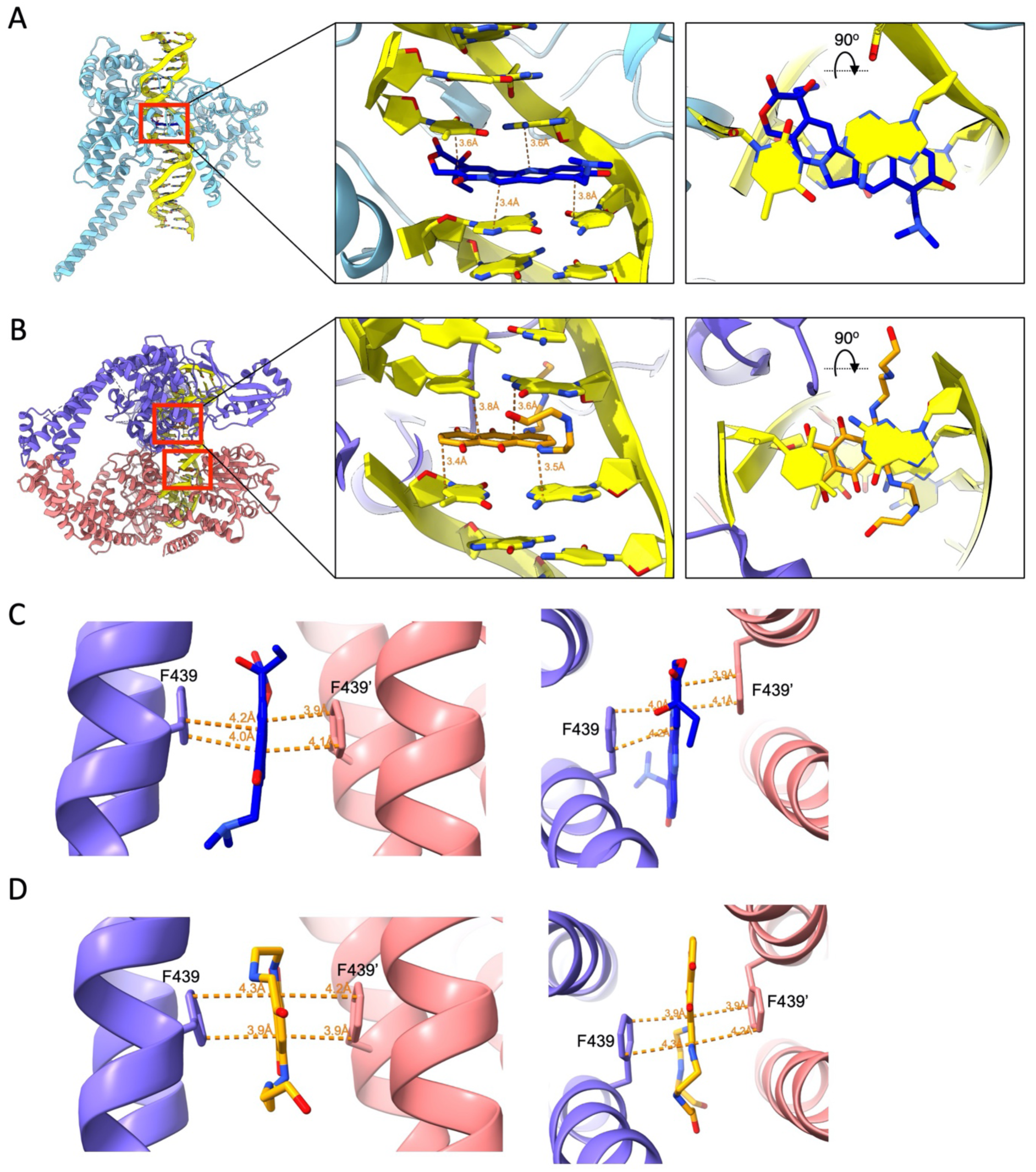
Interactions of topotecan and mitoxantrone with topoisomerases and ABCG2. (**A**), Model of human topoisomerase I in a complex with topotecan and DNA (**<u>PDB 1K4T</u>**). The topoisomerase I is light blue, DNA yellow and topotecan blue. The region with inhibitor was marked with red rectangle and zoomed in to show both side and top views. The distances between aromatic rings of inhibitor and base pairs of nucleotides are shown as dashed lines and colored in orange. (**B**), Model of human topoisomerase II beta in a complex with two mitoxantrones and DNA (**<u>PDB 4G0V</u>**). Two monomers of topoisomerase II were colored purple and salmon, DNA is yellow and the two mitoxantrone molecules orange. Regions with mitoxantrone bound are marked with red rectangles and show as in (**A)**. (**C**), Distances between topotecan and F439s in the binding pocket of ABCG2 transporter. ABCG2 monomers are colored salmon and purple. (**D**), Distances between mitoxantrone and F439s in ABCG2. Colored as in (**C**).

## SUPPLEMENTARY MOVIES

**Movie S1**

Visualization of the 3D variability analysis of ABCG2-topotecan-Fab structure performed in CryoSPARC2. The nanodisc density was masked.

**Movie S2**

Visualization of the 3D variability analysis of ABCG2-mitoxantrone-Fab structure performed in CryoSPARC2. The nanodisc density was masked.

**Movie S3**

Visualization of the 3D variability analysis of ABCG2-tariquidar-Fab structure performed in CryoSPARC2. The nanodisc density was masked.

## Notes

### Competing Interest Statement

The authors have declared no competing interest.

